# Nuclear phosphatidylinositol 3,4,5-trisphosphate interactome uncovers an enrichment in nucleolar proteins

**DOI:** 10.1101/2020.05.17.100446

**Authors:** Fatemeh Mazloumi Gavgani, Andrea Papdiné Morovicz, Clive S. D’Santos, Aurélia E. Lewis

## Abstract

Polyphosphoinositides (PPIn) play essential functions as lipid signalling molecules and many of their functions have been elucidated in the cytoplasm. However, PPIn are also intranuclear where they contribute to chromatin remodelling, transcription and mRNA splicing. The PPIn, phosphatidylinositol 3,4,5-trisphosphate (PtdIns(3,4,5)*P*_3_) has been mapped to the nucleus and nucleoli but its role remains unclear in this subcellular compartment. To gain further insights into the nuclear functions of PtdIns(3,4,5)*P*_3_, we applied a previously developed quantitative mass spectrometry-based approach to identify the targets of PtdIns(3,4,5)*P*_3_ from isolated nuclei. We identified 179 potential PtdIns(3,4,5)*P*_3_-interacting proteins and gene ontology analysis for the biological functions of this dataset revealed an enrichment in RNA processing/splicing, cytokinesis, protein folding and DNA repair. Interestingly, about half of these interactors were common to nucleolar protein datasets, some of which had dual functions in rRNA transcription and DNA repair, including Poly(ADP-Ribose) Polymerase 1 (PARP1/ARTD1). PARP1 was found to interact directly with PtdIns(3,4,5)*P*_3_ as well as PtdIns(3,4)*P*_2_ and to co-localise with PtdIns(3,4,5)*P*_3_ in the nucleolus and with PtdIns(3,4)*P*_2_ in nucleoplasmic foci. In conclusion, the PtdIns(3,4,5)*P*_3_ interactome reported here identified several nucleolar proteins and further pointed to roles for this lipid in these processes.

## Introduction

Polyphosphoinositides (PPIn, nomenclature from (1)) are phosphorylated derivatives of the glycerophospholipid, phosphatidylinositol (PtdIns) (2). The inositol ring can be reversibly phosphorylated at the 3’, 4’ and 5’ hydroxyl groups, producing seven different PPIn, *i.e.* PtdIns3*P*, PtdIns4*P* and PtdIns5*P*, PtdIns(3,4)*P*2, PtdIns(3,5)*P*_2_, PtdIns(4,5)*P*_2_, and PtdIns(3,4,5)*P*_3_ (3). These lipids can act directly as signalling molecules or indirectly as precursors of second messengers. They are metabolised in different subcellular compartments due to the presence of substrate specific PPIn metabolizing kinases and phosphatases (4, 5). While the roles and regulation of PPIn have been extensively studied in the cytoplasm, the importance of their nuclear roles are only recently becoming more apparent (6, 7). The presence of PPIn as well as specific PPIn enzymes was first demonstrated in an intra-nuclear pool not associated with the nuclear envelop (8, 9). The concept of PPIn metabolism and signalling occurring in the nucleus independently of the cytoplasm was reported shortly after in several studies (10–12). Consequently, with the exception of PtdIns(3,5)*P*_2_, the remaining six PPIn have been detected and/or quantified in the nucleus (13–25). The intra-nuclear physico-chemical state of PPIn is still unclear, but several possibilities are emerging to explain how the acyl chains can be shielded from the aqueous environment. These have been shown to be buried in the hydrophobic ligand pocket of the nuclear receptors Liver Receptor Homolog-1 and Steroidogenic Factor 1 while the inositol headgroup remains accessible for modification by PPIn enzymes (26–28). Alternatively, the presence of nuclear lipid droplets has recently been reported in a few studies (29, 30) including the newly discovered nuclear lipid islets, which consist of PtdIns(4,5)*P*_2_ nuclear aggregates possibly in the form of micelles, hence accommodating the acyl chains facing inwards (31).

Several studies have identified multiple nuclear processes attributed to nuclear PPIn, including mRNA processing, splicing and export, chromatin remodelling, transcription as well as cell cycle progression (32–38). Nuclear PPIn regulate these processes by interacting electrostatically with proteins via plekstrin homology (PH) domain in few cases (39, 40) but mostly via polybasic regions (PBR), also called K/R rich motifs ((25, 41–52), and recently reviewed in (7)). So far, PtdIns(4,5)*P*_2_, its metabolising enzymes and effector proteins have been identified in nuclear speckles, hubs of mRNA processing and export (20, 21, 44, 53, 54). Other nuclear PtdIns(4,5)*P*_2_ effector proteins have roles in chromatin remodelling (55, 56), transcriptional regulation and protein stability (48, 57, 58). A minor PtdIns(4,5)*P*_2_ pool was also detected in the nucleolus where it plays a role in RNA polymerase I-mediated transcription (13, 21, 59, 60). Mono-phosphorylated PPIns, interact with several histone-binding proteins (41, 49, 51, 61) or transcription factors or cofactors (43, 45, 50). PtdIns(3,4,5)*P*_3_ and the class I phosphoinositide 3-kinase (PI3K) catalytic subunit p110β are localised in the nucleoplasm and nucleolus (19, 25, 62, 63). Only a few nuclear PtdIns(3,4,5)*P*_3_-effector proteins have so far been reported and include PtdIns(3,4,5)*P*_3_-binding protein (PIP3-BP) (40), the GTPase L-PIKE (L-isoform of PI3K enhancer) (39), the mRNA export protein ALY (THO complex subunit 4) (44), UDP-N-acetylglucosamine-peptide N-acetylglucosaminyltransferase (OGT) (64) as well as the nucleolar proteins nucleophosmin and ErbB3-binding protein 1 (EBP1) (25, 42). Overall, although PtdIns(3,4,5)*P*_3_ is likely to be a key signalling PPIn in the nucleus, its nuclear function remain largely unknown.

To identify proteins interacting specifically with PtdIns(3,4,5)*P*_3_, several interactomics studies have been performed from a variety of cell types using either cytosolic (65–67) or whole cell extract (68). To enrich for nuclear PPIn-interacting proteins, for which the least is known, we developed a PPIn quantitative interactomics approach using isolated nuclei and based on an enrichment of PPIn interactors (46). This approach led to the identification of PtdIns(4,5)*P*_2_ nuclear interacting partners involved in mRNA transcription regulation, mRNA splicing and protein folding. In this study, we have performed quantitative mass spectrometry-based PtdIns(3,4,5)*P*_3_ interactomics from isolated HeLa nuclei using the same approach (46). We identified 179 potential PtdIns(3,4,5)*P*_3_ interactors with functions highly enriched in protein folding, RNA splicing, DNA repair and cell cycle regulation. Interestingly, half of these proteins were common to the T cell nucleolome protein list (69). In this study, we focused on Poly(ADP-Ribose) Polymerase 1 (PARP1, now referred as ADP-ribosyltransferase 1, ARTD1), validated its direct interaction with PtdIns(3,4,5)*P*_3_ and showed its colocalisation in nucleoli. In sum, this study validates our approach to identify globally PPIn-interacting proteins based in the nucleus and represents a resource for further research efforts investigating the role of PtdIns(3,4,5)*P*_3_ in these interactions.

## Results

### PtdIns(3,4,5)*P*_3_ is nucleolar in HeLa cells

To extend our previous findings on the nucleolar localisation of PtdIns(3,4,5)*P*_3_ previously observed in the breast cancer cell line AU565 (25), we determined its subcellular localization in actively growing HeLa cells by immunofluorescence staining and confocal microscopy (Figure 1). Using specific antibodies to detect PtdIns(3,4,5)*P*_3_ ((25) and Figure 1B), we observed the presence of this PPIn in the nucleolus in 74% +/− 10% of asynchronous HeLa cells in either intense or diffuse foci which colocalised with the nucleolar proteins nucleolin or the transcription factor upstream binding factor (UBF) (Figure 1D and supplementary Figure S1A). In addition, the presence of PtdIns(3,4,5)*P*_3_ in the nucleolus was supported using the purified PH domain of the general receptor of phosphoinositides-1 (GRP1, alias cytohesin-3) conjugated to EGFP and GST as a labelling probe. The PH domain of GRP1 is well known for its affinity to PtdIns(3,4,5)*P*_3_ while the K273A point mutation disrupts this interaction (70–73). When tested by lipid overlay assay, the WT GST-EGFP-GRP1-PH demonstrated interaction with PtdIns(3,4,5)*P*_3_ but not the K273A mutant (Figure 1C). Labelling with this probe highlighted foci within rings detected by the nucleolar protein nucleophosmin when using the WT protein in about 50% of cells (Figure 1E-F). In contrast, the percentage of cells showing these foci was greatly reduced to 6% when using the K273A mutant (Figure 1E-F).

**Figure 1.**
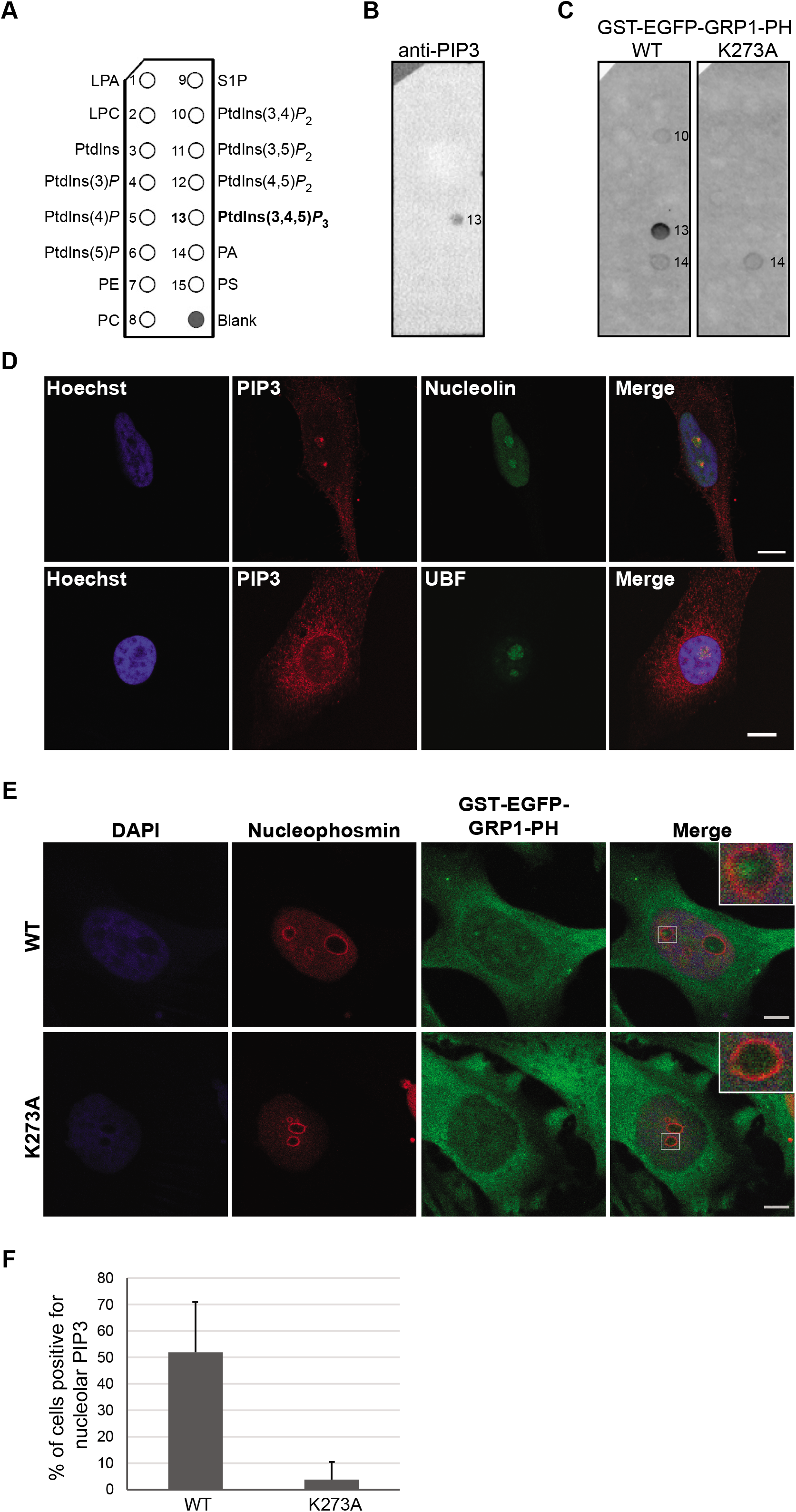
Specific detection of PtdIns(3,4,5)*P*_3_ in the nucleolus. **(A)** PIP strips (Echelon Inc) schematic showing the position of the spotted lipids each with 100 pmol. LPA, lysophosphatidic acid; LPC, lysophosphatidylcholine; PI, phosphatidylinositol; PE, phosphatidylethanolamine; PC, phosphatidylcholine; S1P, sphingosine-1-phosphate; PA, phosphatidic acid; PS, phosphatidylserine. **(B)** Validation of anti-PtdIns(3,4,5)*P*_3_ antibody specificity using PIP strips. **(C)** Validation of the specificity of the recombinant GST-EGFP-GRP1-PH WT versus binding mutant K273A. **(D, E)** Confocal microscopy of actively growing HeLa cells stained with the indicated antibodies (D) or by incubation with recombinant GST-EGFP-GRP1-PH WT or K273A mutant combined with anti-nucleophosmin staining (E). **(F)** Quantification of the detection of nucleolar PtdIns(3,4,5)*P*_3_ expressed as the percentage of HeLa cells + SDs showing foci detected by the GST-EGFP-GRP1-PH probe WT or K273A mutant within the area delimited by nucleophosmin. PIP3: PtdIns(3,4,5)*P*_3_; UBF: upstream binding factor; GRP1: general receptor for phosphoinositides-1; PH: plextrin homology. Scale bar represents 5 μm.

### The nuclear Ptdlns(3,4,5)*P*_3_ interactome is enriched in nucleolar proteins

The existence of PtdIns(3,4,5)*P*_3_ in the nucleus has been reported (19, 24, 25, 63), but so far only a few nuclear proteins have been reported to interact with PtdIns(3,4,5)*P*_3_ and knowledge of its function is limited in this cell compartment. We sought to identify the interacting partners of PtdIns(3,4,5)*P*_3_ in the nucleus using a quantitative proteomics method previously developed for the identification of nuclear PtdIns(4,5)*P*_2_ effector proteins (46) with a view to identifying nuclear processes that PtdIns(3,4,5)*P*_3_ may regulate,. Following SILAC labelling of HeLa S3 cells, nuclei were isolated and incubated with neomycin to enrich for and displace potential PPIn-binding proteins from nuclei (Figure 2A). Equal protein amounts obtained from heavy labelled and light labelled cell populations were incubated with PtdIns(3,4,5)*P*_3_-conjugated beads or control beads respectively. The specificity of the PtdIns(3,4,5)*P*_3_ affinity beads was validated by pull down assay with GST-GRP1-PH (Figure 2B). The control beads showed no affinity whereas the PtdIns(3,4,5)*P*_3_ beads were able to pull down the GST-GRP1-PH domain. Importantly, this interaction was negligible in the preincubation of free PtdIns(3,4,5)*P*_3_ with the probe. The pull down eluates were combined and separated by SDS-polyacrylamide gel electrophoresis (PAGE). Following trypsin digestion, the peptides were analysed by LC-MS/MS and identified and quantified using Sequest search engine. Statistical analyses demonstrated 179 proteins with at least two peptides to be specifically pulled down by PtdIns(3,4,5)*P*_3_, 75 (42%) of which were identified in two additional experiments (Figure 2C and Supplementary Table S1). These included proteins previously reported experimentally as *bona fide* nuclear PtdIns(3,4,5)*P*_3_ interacting proteins, *i.e.* nucleophosmin (42), ALY (THO complex subunit 4) (44), IQ motif containing GTPase-activating proteins (IQGAP1) (74) as well as OGT (64). In addition to these proteins, 20 from our dataset had previously been identified in PtdIns(3,4,5)*P*_3_ interactomes from whole cell extracts (67, 68) (Figure 2D and Table 1). Several proteins were also common to the nuclear PtdIns(4,5)*P*_2_ interactome ((46), Table 2). Importantly, the majority of the identified proteins are likely to be direct PtdIns(3,4,5)*P*_3_ interactions, since only a few clusters involved in proteinprotein complexes were detected using the STRING web tool (Supplementary Figure S2). We further searched for the presence of PPIn-binding domains and found only 4 proteins, including dynamin 1, 2 and 3 harbouring a PH domain with previous knowledge of PPIn interaction (75, 76)) or ATP binding cassette sub family F member 1 with the less studied PDZ domain (77). In contrast, the lysine/arginine rich motif (K/R-(X_n=3-7_)-K-X-K/R-K/R), which we previously reported to be enriched in PtdIns(4,5)*P*_2_-binding proteins (46), was found in 38% of PtdIns(3,4,5)*P*_3_-associated proteins, accounting for a 1.4 fold enrichment compared to proteins pulled down by control beads (Figure 2C and Supplementary Table S1) and 1.3 fold compared to proteins annotated to the nucleus (nucleome). For a better understanding of the biological processes of these proteins, they were mapped to the Gene Ontology (GO) database for biological processes and an enrichment test was performed using the PANTHER 14.1 web tool (2019-03-12 release, (78, 79)). The biological processes that were over-represented by >5 fold are shown in Figure 2E. In particular, RNA splicing/processing, protein folding, cytokinesis and DNA repair were functions particularly enriched in the PtdIns(3,4,5)*P*_3_ pull down protein list. A large number of the potential PtdIns(3,4,5)*P*_3_ interactors were linked or annotated to the nucleolus, as highlighted in Supplementary Table S1. Indeed, 17% of all potential PtdIns(3,4,5)*P*_3_ interactors are common to the nucleolar database (80) and 51% to the T cell nucleome (69), including 19 common to both nucleome lists.

**Figure 2.**
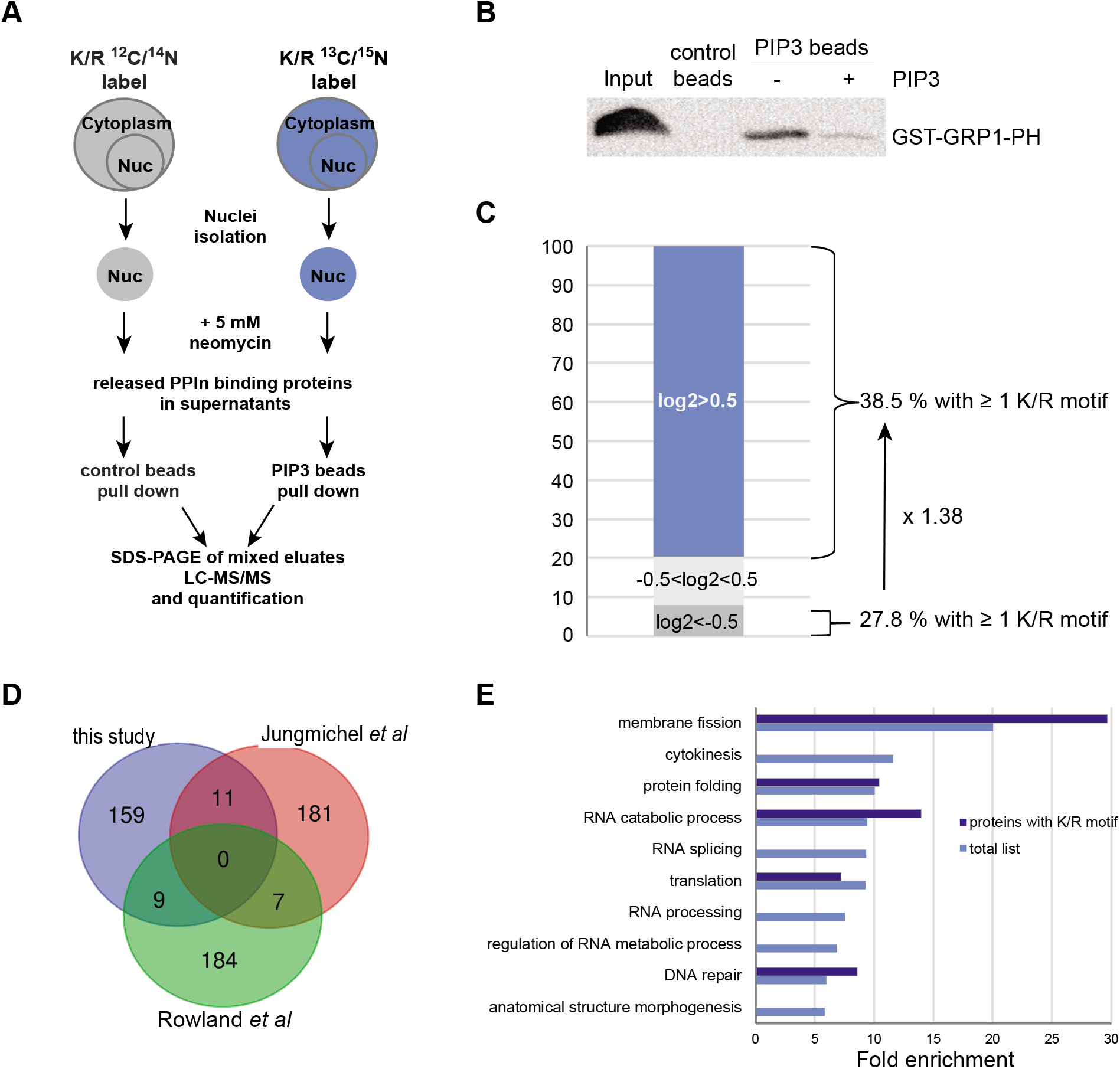
Nuclear PtdIns(3,4,5)*P*_3_ interactome. **(A)** Workflow of the experimental setup where heavy labelled and unlabelled HeLa S3 nuclei were incubated with 5 mM neomycin and the displaced proteins pulled down using control beads or PtdIns(3,4,5)*P*_3_-conjugated beads and subsequently analysed by LC-MS/MS. **(B)** GST-GRP1-PH pull down with control or PtdIns(3,4,5)*P*_3_-conjugated beads in the absence (-) or presence (+) of 20 μM free PtdIns(3,4,5)*P*_3_ (PIP3). Eluates were resolved by SDS-PAGE and western immunoblotted using an anti-GST antibody conjugated to horse radish peroxidase. **(C)** Protein distribution in % according to their Log2 values: proteins with Log2< −0.5 (binding to control beads, 18 proteins), −0.5<Log2<0.5 (proteins binding equally to PtdIns(3,4,5)*P*_3_ or control beads, 26 proteins) and Log2>0.5 (binding to PtdIns(3,4,5)*P*_3_ beads, 179 proteins). **(D)** Venn diagram comparing the PtdIns(3,4,5)*P*_3_-interactome from this study to two others (67, 68). **(E)** Biological processes Gene ontology fold enrichment of the proteins pulled down specifically by the PtdIns(3,4,5)*P*_3_-conjugated beads from this study with FDR *P* <0.05 and with at least 5 proteins in each process.

**Table 1.**
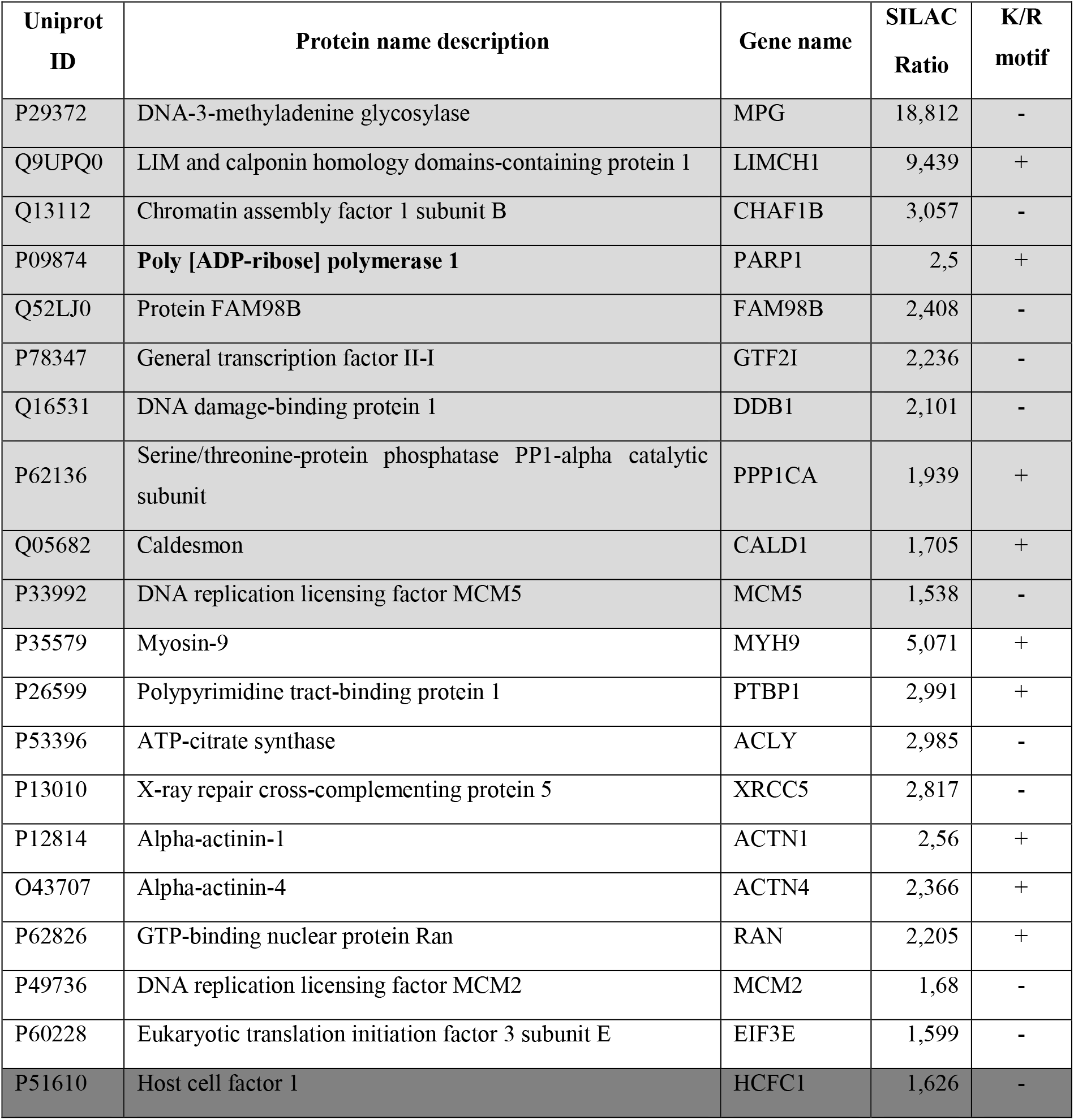
List of PtdIns(3,4,5)*P*_3_ binding proteins common to two other reported PtdIns(3,4,5)*P*_3_ interactomes. Proteins pulled down by PtdIns(3,4,5)*P*_3_ identified in this study common to those in PtdIns(3,4,5)*P*_3_ interactome lists from Jungmichel *et al* (68), indicated in light grey, Rowland *et al* (67) in white and Bidlingmaier *et al* (47) in dark grey.

**Table 2.** List of PtdIns(3,4,5)*P*_3_ binding proteins common to the nuclear PtdIns(4,5)*P*_2_ interactome. Proteins pulled down by PtdIns(3,4,5)*P*_3_ (PIP3) identified in this study common to those reported in the PtdIns(4,5)*P*_2_ (PIP2) nuclear interactome that we have previously published (46). Protein highlighted in bold indicate proteins with roles in splicing.

| Uniprot ID | Protein name description | Gene name | SILAC Ratio |  |
| --- | --- | --- | --- | --- |
|  |  |  | PIP3 list | PIP2 list |
| P11021 | 78 kDa glucose-regulated protein | HSPA5 | 1.787 | 1.63 |
| Q9P258 | Protein RCC2 | RCC2 | 2.305 | 1.869 |
| **P26599** | **Polypyrimidine tract-binding protein 1** | **PTBP1** | **2.991** | **1.964** |
| **O60506** | **Heterogeneous nuclear ribonucleoprotein Q** | **SYNCRIP** | **1.504** | **2.033** |
| P68104 | Elongation factor 1-alpha 1 | EEF1A1 | 2.007 | 2.072 |
| **Q99729** | **Heterogeneous nuclear ribonucleoprotein A/B** | **HNRNPAB** | **2.604** | **2.272** |
| **Q14103** | **Heterogeneous nuclear ribonucleoprotein D** | **HNRNPD** | **2.345** | **2.44** |
| P23284 | Peptidyl-prolyl cis-trans isomerase B | PPIB | 2.342 | 2.450 |
| **Q00839** | **Heterogeneous nuclear ribonucleoprotein U** | **HNRNPU** | **1.742** | **2.676** |
| P26641 | Elongation factor 1-gamma | EEF1G | 1.565 | 2.84 |
| P29692 | Elongation factor 1-delta | EEF1D | 2.078 | 2.89 |

**Table 3.**
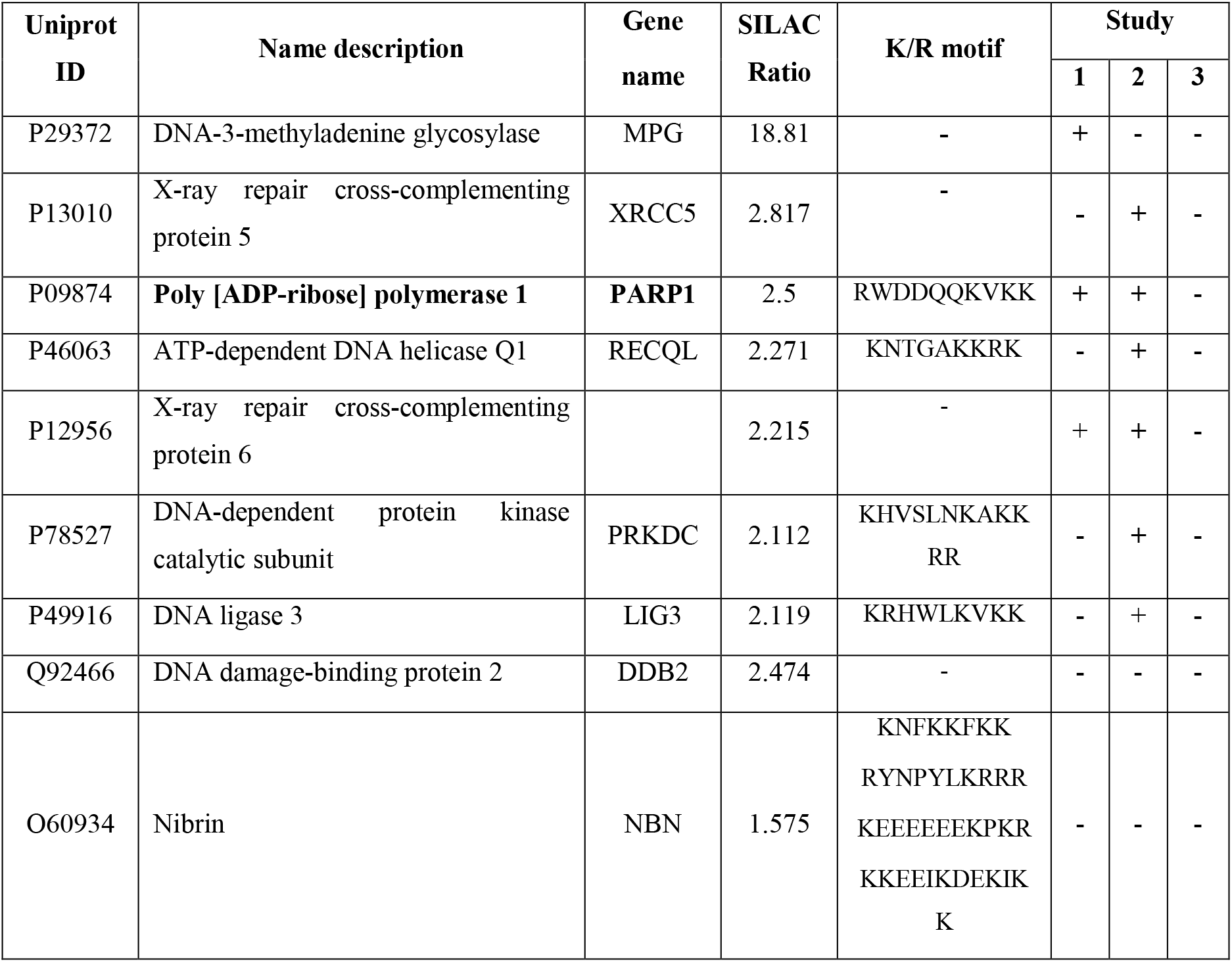
List of potential PtdIns(3,4,5)*P*_3_ binding proteins annotated to DNA repair. Proteins pulled down by PtdIns(3,4,5)*P*_3_ and annotated to the DNA repair enriched process, identified with at least 2 peptides, with heavy/light log2 ratios >0.5, are indicated in this table. Their presence (highlighted +) or absence (-) in the nucleolar database (study 1, (80)), the T cell nucleome (study 2, (69)) and/or the HeLa nucleome (study 3, (92)) is indicated. K/R motifs consists of the following sequence: K/R-(X_n=3-7_)-K-X-K/R-K/R.

### PtdIns(3,4,5)*P*_3_ co-localizes with PARP1 in nucleoli

PARP1, a chromatin-associated protein previously reported to be abundant in the nucleolus (81, 82) and which harbours one K/R motif, was identified as a specific PIP3 interacting protein with a PtdIns(3,4,5)*P*_3_/control SILAC ratio of 2.5 (Table 1 and Supplementary Table S1). We first biochemically validated the direct interaction of PARP1 with PPIn by lipid overlay assay using phospholipid-immobilized strips and GST-PARP1 recombinant protein (Figure 3A). PARP1 was found to interact with most PPIns as well as phosphatidic acid and phosphatidylserine but not with other glycerophospholipids or sphingolipid. In contrast, GST alone showed no interaction. We validated these results using a PPIn pull down assay and demonstrated narrower specificity of interaction of GST-PARP1 to PtdIns(3,4)*P*_2_ and PtdIns(3,4,5)*P*_3_ and lack of interaction to PtdIns(4,5)*P*_2_ or the monophosphorylated PPIn (Figure 3B). To control for the lack of interaction with PtdIns(4,5)*P*_2_, we tested the PtdIns(4,5)*P*_2_-conjugated beads with the PH domain of phospholipase Cδ1 and showed a strong interaction. The nuclear localisation of PtdIns(4,5)*P*_2_ and PtdIns(3,4,5)*P*_3_ is well established and we compared the endogenous localisation of these lipids to PARP1 by immunofluorescent staining. PtdIns(3,4,5)*P*_3_ co-localized with PARP1 in the nucleolus of HeLa cells (Figure 3C and Supplementary Figure S2A). In contrast, PtdIns(4,5)*P*_2_, segregated to nuclear speckles consistently to previous studies and mostly did not colocalise with PARP1. Considering the interaction of PARP1 with PtdIns(3,4)*P*_2_-in the pull down assay, we tested their localisation and showed that nucleoplasmic PARP1 co-localised with PtdIns(3,4)*P*_2_-positive foci (Figure 3C).

**Figure 3.**
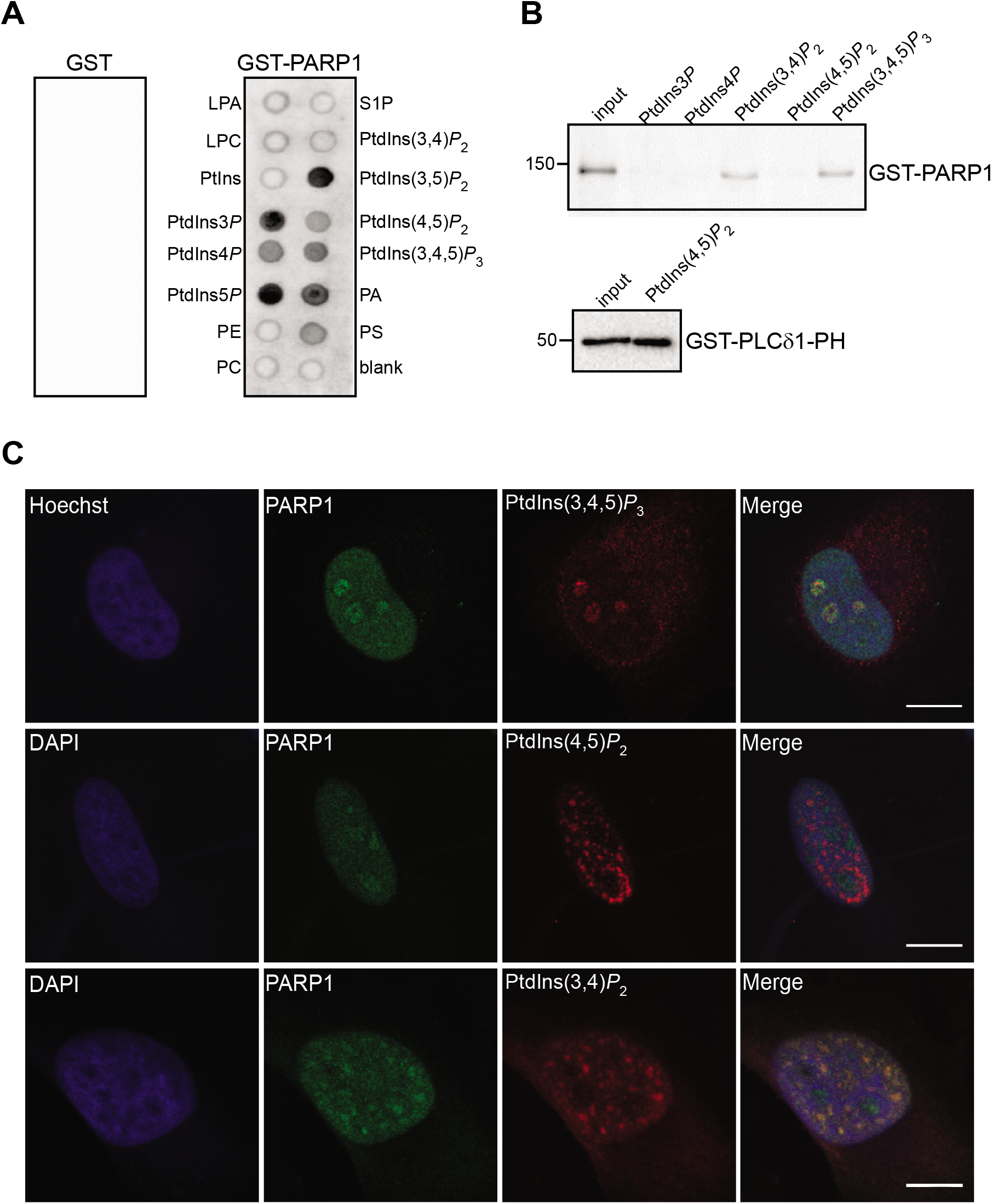
PtdIns(3,4,5)*P*_3_ interacts and colocalises with PARP1 in the nucleolus. **(A)** PIP strips incubated with recombinant GST or GST-PARP1 and detection of protein-lipid interactions using an anti-GST-HRP conjugated antibody. **(B)** GST-PARP1 or GST-PLCδ1 pull down with the indicated PPIn-conjugated beads. Eluates were resolved by SDS-PAGE and western immunoblotted using an anti-GST antibody conjugated to horse radish peroxidase. **(C)** HeLa cells costained with anti-PARP1 and PtdIns(3,4,5)*P*_3_ antibodies and imaged by confocal microscopy. Scale bar represents 20 μm in the top image and 10 μm in the bottom image.

## Discussion

Evidence of the presence of PPIn in the nucleus together with the kinases responsible for their synthesis is now well established (34, 38, 83–85). Interestingly, they are found in RNA-rich membrane-less compartments, such as the nuclear speckles and nucleolus in particular for PtdIns(4,5)*P*_2_ (20–22, 53) and PtdIns(3,4,5)*P*_3_ (25) respectively. In this study, we have extended our previous findings (25) by showing the localization of PtdIns(3,4,5)*P*_3_ in the nucleolus in HeLa cells. To support these findings, a minor pool of PtdIns(4,5)*P*_2_ has been previously reported in the nucleolus and could hence substantiate the nucleolar synthesis of PtdIns(3,4,5)*P*_3_ (21, 59). In addition, the PPIn kinase isoforms, PI4K IIα, PIP5K Iα and PI3K p110β which synthesise PtdIns 4*P*, PtdIns(4,5)*P*_2_ and PtdIns(3,4,5)*P*_3_ respectively, have all been shown to be present in the nucleolus (25, 86, 87). Similarly, evidence exists for PtdIns(3,4,5)*P*_3_ phosphatases PTEN (phosphatase and tensin homolog) and SHIP1 (Src homology 2-domain-containing inositol phosphatase) in the nucleolus (88, 89). All the components are therefore in place in nucleoli for the regulation of PtdIns(3,4,5)*P*_3_ synthesis and a potential role in this sub-nuclear compartment. The physico-chemical form in which PtdIns(3,4,5)*P*_3_ exist in a non-membranous environment such as the nucleolus is unclear. How the acyl chains can be sheltered from the aqueous environment may be explained by the formation of micelles from the aggregation of acyl chains (83, 90) but this has not been demonstrated so far. Aggregates of PtdIns(4,5)*P*_2_ in nuclear lipid islets (NLIs) in the nucleoplasm have recently been described and consist of proteo-lipid aggregates of about 100 nm in size (31). The PtdIns(3,4,5)*P*_3_ foci detected via confocal microscopy may indicate the presence of PtdIns(3,4,5)*P*_3_ nucleolar aggregates in the form of islets consistent with the PtdIns(4,5)*P*_2_-associated NLIs. This would imply that the acyl chains of PtdIns(3,4,5)*P*_3_ are shielded from the nuclear environment within the core of micelle-like foci, giving a plausible explanation for the biophysical presence of such lipids in the absence of membrane. This remains however to be explored.

To further decipher the role(s) of nuclear PtdIns(3,4,5)*P*_3_, we applied a quantitative interactomics method that we had previously developed to identify PtdIns(4,5)*P*_2_ nuclear effectors (46). To this end, we have identified 179 proteins specifically pulled down by PtdIns(3,4,5)*P*_3_ and not by control beads. Our study allowed the identification of nuclear effector proteins, the majority of which were not identified in the previous interactome performed from whole cell extracts (68). The highest proportion of PtdIns(3,4,5)*P*_3_ interactors were indeed annotated to other compartments than the nucleus in that study, hence masking potential nuclear effector proteins. Except for a few proteins known to be engaged in proteinprotein complexes, the remaining proteins identified in this study are likely to be direct PtdIns(3,4,5)*P*_3_-interactors. Several proteins have previous history as nuclear PPIn effector proteins, such as nucleophosmin, OGT, IQGAP1 and ALY and have well characterised PPIn-binding sites (42, 44, 64, 74), hence validating our approach. When searching for the presence of PPIn domains, only three proteins identified in this study have structured PPIn-interacting domains, *i.e.* dynamin 1-3, each harbouring a PH domain (75, 76). The PH domain of these proteins has been reported to bind to PtdIns(4,5)*P*_2_ (75, 76) but also to PtdIns(3,4,5)*P*_3_ in another study (91). Although dynamin family members are GTPase considered to be localised on membranes and microtubules, dynamin-2 and −3 were identified in nucleolar proteomes (69, 92). The function of dynamin in the nucleolus has not been investigated so far. IQGAP1 binds to PtdIns(3,4,5)*P*_3_ via an atypical PPIn binding domain lined with basic residues, with a distinct fold to most known domains (74). Although IQGAP1 has clear roles in the cytoplasm, it was also reported to accumulate in the nucleus at the G1/S phase of the cell cycle (93). Still how PPIn binding affects its nuclear role is unknown.

The majority of the PtdIns(3,4,5)*P*_3_-interactors identified in this study are characterised by the presence of at least one polybasic motif shown previously to serve as PPIn interaction sites via electrostatic interactions in other nuclear proteins (25, 41, 43, 45, 48–52) or of basic patches, as found in nucleophosmin, ALY and OGT (42, 44, 64). This finding is consistent with the enrichment of such motifs in the nuclear PtdIns(4,5)*P*_2_ interactome that we have previously reported (46). These findings would point to a predominant mode of interaction via short polybasic motifs in membrane-less compartments of the nucleus compared to cytoplasmic membrane-mediated interaction via PPIn-binding domains.

The PtdIns(3,4,5)*P*_3_-binding protein list was highly enriched in nucleolar proteins, including nucleophosmin. The nucleolus is a compartment where rRNA transcription and processing occur to enable ribosome biogenesis (94). However, gene ontology analyses of nucleolar proteomes showed their association with other biological functions such as cell cycle regulation and DNA repair (94, 95). Indeed, a growing body of evidence indicates that some nucleolar proteins have roles in DNA repair (95–98). Interestingly, among the PtdIns(3,4,5)*P*_3_ interacting proteins identified in this study, an enrichment of DNA repair proteins was shown, listing nine proteins, seven of which were found in at least one of the nucleolome datasets previously published, including PARP1. PARP1 has also recently been identified in the PPIn interactome from whole cell extracts reported by Jungmichel *et al* showing specificity for PtdIns(3,4,5)*P*_3_ (68). In this study, we have shown that PARP1 binds directly to PPIn *in vitro* using lipid overlay assay and PPIn pull down. The pull down assay showed some interaction specificity towards PtdIns(3,4,5)*P*_3_ and PtdIns(3,4)*P*_2_ compared to the lipid overlay assay showing little specificity for the different PPIn species. Lipid presentation is different in these two assays and include the whole molecule when the PPIn-conjugated beads are used. This may explain the difference in specificity and may suggest that hydrophobic interactions contribute to the specificity of interaction. Consistent with *in vitro* interaction studies, PARP1 co-localised with PtdIns(3,4,5)*P*_3_ in the nucleolus and with PtdIns(3,4)*P*_2_ in nucleoplasmic foci, and not with PtdIns(4,5)*P*_2_. The localisation of PtdIns(3,4)*P*_2_ in nucleoplasmic foci is consistent with a recent study by Kalasova *et al* (13). The identity of these foci has not been investigated but appear to be distinct to PtdIns(4,5)*P*_2_-positive sites which localises to nuclear speckles (21) but not with PARP1. Knowledge of the synthesis route of this PPIn in the nucleus is limited but was shown in one study to be produced by the 5-phosphatase, SHIP2, by dephosphorylating PtdIns(3,4,5)*P*_3_ in vascular smooth muscle cells (99). SHIP2 (99) or in its phosphorylated form on serine 132 (100) was found in nuclear speckles in different cells. Alternatively, the class II PI3K, PI3KC2α, known to produce PtdIns(3,4)*P*_2_ by phosphorylating PtdIns4*P*, was also reported to localise in nuclear foci (101).

PARP1 does not harbour any known PPIn-binding domain but has a K/R motif in the first zinc finger domain as well as polybasic regions in the two other zinc finger domains. These basic regions could provide a mechanism of interaction to PPIn consistently with other nuclear proteins found previously to interact with PPIn (7). Lack of specificity *in vitro* consistent with other nuclear proteins with the same reported mode of interactions.

The nucleolar protein, nucleophosmin has previously been shown to bind the DNA binding domain of PARP-1 (82) and it is in addition a well-known PtdIns(3,4,5)*P*_3_ interacting protein (42). When cells are not under stress conditions, an enrichment of both PARP1 and poly ADP-ribose can be observed in the nucleolus (82, 102). Upon RNA polymerase I inhibition, PARP1 delocalizes from the nucleolus, indicating that the presence of PARP1 in the nucleolus is dynamic and dependent on RNA polymerase I transcriptional activity. Nucleolar delocalization of PARP1 is accompanied by other nucleolar proteins such as nucleophosmin and UBF (82, 103, 104). Altogether, these studies suggest that the organisation of proteins and lipids within the nucleolus is affected by the active transcription of rRNA. Interestingly, both PARP1 and nucleophosmin are also histone H1 interacting proteins (105), which is emerging to play important roles in the structure and integrity of the nucleolus (106). PARP1-dependent PARylation of histone H1 has been shown to remove this histone from the chromatin, hence causing it to relax (107). Nucleophosmin binds to histone H1.5 and has a silencing effect on this linker histone (105). A link between histone H1 and PPIn has previously been reported, demonstrating its interaction with PtdIns(4,5)*P*_2_ via its C-terminal region (57). Although Yu *et al* did not study the possible interaction of histone with PtdIns(3,4,5)*P*_3_, this PPIn may form a complex with PARP1, nucleophosmin and H1 to regulate the architecture of the nucleolus, which may allow transcription to occur.

In conclusion, this study extends our knowledge of PtdIns(3,4,5)*P*_3_ interacting proteins already identified from cytoplasmic or whole cell extract sources and further acknowledge the complexity of these interactions in the nucleus. Our approach based on neomycin-dependent displacement of proteins allowed the identification of numerous nuclear PtdIns(3,4,5)*P*_3_ binders. This resource is amenable for further biochemical and functional characterisation assessing how the array of nuclear, and in particular nucleolar, functions these interactions can regulate.

## Materials and methods

### Cell culture and SILAC labelling

HeLa cells were grown in DMEM medium containing 10% fetal bovine serum (FBS) in 5% CO_2_ at 37°C. For SILAC (Stable isotope labelling with amino acids in cell culture) labelling, HeLa S3 cells were grown in heavy (^13^C6,^15^N_2_-labelled lysine and ^13^C_6_, ^15^N_4_-labelled arginine) or light (unlabelled amino acids) DMEM medium (Silantes, cat# 280001300) supplemented with 10% dialyzed FBS (Silantes, cat# 281000900). To examine the efficiency of SILAC labelling the incorporation of heavy amino acids was validated by LC-MS by Dr Bernd Thiede (University of Oslo, Norway).

### Nuclear fractionation

Cells were grown in 10 x 15 cm dishes up to 70% confluency. 1 h after adding fresh medium, the cells were washed, trypsinized and washed again 3 times (this time with ice cold PBS). The cell pellet was re-suspended in 5 ml of buffer A (10 mM HEPES pH 7.9, 1.5 mM MgCl_2_, 10 mM KCl, 0.5 mM DDT, 1% Igepal and protease inhibitor cocktail) and incubated on ice for 5 min. The cells were then passed 12 times through a 23-gauge needle to disrupt the cell membrane. The lysates were then centrifuged at 218x g for 5 min at 4°C. The supernatant was collected as the cytosolic fraction and the pellet containing the nuclei was re-suspended in 3 ml of buffer S1 (0.25 M sucrose, 10 mM MgCl_2_ and protease inhibitor cocktail) and layered over 3 ml of buffer S2 (0.35 M sucrose, 0.5 mM MgCl_2_ and protease inhibitor cocktail) and centrifuged at 1430xg for 5 min at 4 °C. Crude nuclear fractionation was performed as previously reported (25).

### Neomycin extraction

Nuclei were washed with retention buffer containing 20 mM Tris pH 7.5, 70 mM NaCl, 20 mM KCl, 5 mM MgCl_2_, 3 mM CaCl_2_ and protease inhibitor cocktail. The nuclei were then incubated with freshly prepared 5 mM neomycin (Neomycin trisulfate salt, Sigma-Aldrich) in retention buffer, rotating for 30 min at RT. After centrifugation at 13000 rpm for 5 min, the supernatant containing the neomycin-displaced protein extract was collected. Neomycin supernatants were dialysed three times in 900 ml of cold lipid pulldown buffer containing 20 mM HEPES pH 7.5, 150 mM NaCl, 5 mM EDTA, 0.1 % Igepal using Slide-A-Lyser Mini dialysis units (Thermo Fisher) for 1 h at 4°C each time. The protein concentration was measured using BCA (bicinchoninic acid) protein assay (ThermoFisher Scientific).

### PPIn pull down

#### PtdIns(3,4,5)P_3_ pull down for MS

Equal amount of dialysed neomycin supernatants were used for each pulldown. The heavy extracts were incubated with 100 μl PtdIns(3,4,5)*P*_3_-conjugated bead slurry (Echelon Biosciences P-B345a) and the light extracts were incubated with control beads (Echelon Biosciences P-B000) for 1 h rotating at 4°C. The beads were then washed 3x with lipid pulldown buffer (20 mM HEPES pH 7.5, 150 mM NaCl, 5 mM EDTA, 0.1% Igepal) containing phosphatase inhibitors (5 mM β-glycerophosphate, 5 mM NaF and 2 mM Na_3_VO_4_) and protease inhibitor cocktail. The beads were eluted into 2 equal samples and run on SDS-PAGE through the stacking gel only and proteins were stained with Coomassie blue.

#### In vitro PPIn pull down

To test the interaction capacity of PPIn-conjugated beads, lipid pull downs were performed using 1-2 μg GST-GRP1-PH or PLCδ1-PH (expressed and purified as described in (46)) and 10 μl of PtdIns(3,4,5)*P*_3_- and PtdIns(4,5)*P*_2_ -conjugated beads slurry (Echelon Biosciences) respectively. 20 μM PtdIns(3,4,5)P_3_ diC8 (Echelon P-3908) was used in pre incubation. For the PARP1 PPIn pull down, 1.5 μg GST-PARP1 was used together with 15 μl PPIn-conjugated bead slurry.

### Proteomics

#### In-gel digestion

In-gel trypsin digestion was performed as described (108) with some modifications. Briefly, the Coomassie brilliant blue-stained protein bands were excised, and following several washes, the gel pieces were subjected to a reduction step using 10 mM DTT in 100 mM ammonium bicarbonate (NH_4_HCO_3_) buffer for 45 min at 56°C. Alkylation was performed with 55 mM iodoacetamide in 100 mM NH_4_HCO_3_ for 30 min at room temperature in the dark. Digestion was performed with 10 μl of trypsin (10 mg/l in 50 mM NH_4_HCO_3_) overnight at 37^0^C. Eluted peptides were recovered, and the gel pieces were subsequently washed in 2.5% formic acid/80% acetonitrile for 30 min at 37^0^C. The acid wash was combined with the original peptide eluate and dried. Samples were resuspended in 0.1% formic acid and analysed directly by nano-LC-MS/MS.

#### Nano LC-MSMS

Digested peptide mixtures were analysed by nano-LC-MS/MS. Mass spectrometry (MS) was performed using a QExactive HF (Thermo Scientific) coupled to an Ultimate RSLCnano-LC system (Dionex). Optimal separation conditions resulting in maximal peptide coverage were achieved using an Acclaim PepMap 100 column (C18, 3 μm, 100 Å) (Dionex) with an internal diameter of 75 μm and capillary length of 25 cm. A flow rate of 300 nl/min was used with a solvent gradient of 5% B to 45% B in 85 min followed by increasing the gradient to 95% B over 5 min. Solvent A was 0.1% (v/v) formic acid, 5%DMSO in water, whereas the composition of solvent B was 80% (v/v) acetonitrile, 0.1% (v/v) formic acid, 5% DMSO in water.

The mass spectrometer was operated in positive ion mode using an N^th^ order double-play method to automatically switch between Full scan acquisition of peptide precursor ions and HCD generated fragments both using the Orbitrap mass analyser. Survey full-scan MS spectra (from 400 to 1,600 m/z) were acquired in the Orbitrap with resolution (R) 60,000 at 400 m/z (after accumulation to a target of 3,000,000 charges). The method used allowed sequential isolation of the 10 most intense ions for fragmentation, depending on signal intensity, using HCD at a target value of 20,000 charges and resolution of 30,000. Target ions already selected for MS/MS were dynamically excluded for 30 s. Unassigned and 1+ charges were excluded from fragmentation selection. General MS conditions were electrospray voltage, 2.5 kV with no sheath or auxiliary gas flow, an ion selection threshold of 2,000 counts for MS/MS, an activation Q value of 0.25, activation time of 12 ms, capillary temperature of 200°C, and an S-Lens RF level of 60% were also applied. Charge state screening was enabled, and precursors with unknown charge state or a charge state of 1 were excluded. Raw MS data files were processed using Proteome Discoverer v.2.1 (Thermo Scientific). Processed files were searched against the SwissProt human database using the Mascot search engine version 2.3.0. Searches were done with tryptic specificity allowing up to one missed cleavage and a tolerance on mass measurement of 10 ppm in MS mode and 20 ppm for MS/MS ions. Structure modifications allowed were oxidized methionine, and deamidation of asparagine and glutamine residues, which were searched as variable modifications. Using a reversed decoy database, false discovery rate (FDR) was less than 1%. Only proteins identified with at least 2 peptides (and including at least 1 unique peptide) and common to the two replicate runs were kept.

### Bioinformatic analyses

For the K/R polybasic motifs search, an in lab Linux shell script was used to first download the sequences of the PtdIns(3,4,5)*P*_3_ pulled down proteins from Uniprot (curl https://www.uniprot.org/uniprot/) (using the curl tool) and search for the (K/R-(X_3-7_)-K-X-K/R-K/R) motif was then carried out using the grep tool.

For the enrichment analyses, the identified Uniprot entries were statistically compared to those of the human genome restricted to entries annotated to the nucleus compartment (GO:0050789) using PANTHER classification system version 13.1 (78, 79). The representation for each GO category for biological processes was calculated as the ratio between the cluster (PtdIns(3,4,5)*P*_3_ dataset) frequency and the reference dataset (human nucleome) frequency, the frequency being the percentage of gene entries in a particular GO term category compared with the respective total number of entries. Only enriched categories with p values <0.05 are presented.

The presence of structured PPIn domain was assessed via the SMART batch search (http://smart.embl-heidelberg.de/smart/batch.pl).

STRING analysis of all PtdIns(3,4,5)*P*_3_-binding protein entries was based upon experimental prediction methods and a confidence score > 0.9.

### Immunofluorescence staining and microscopy

HeLa cells grown on 12 mm coverslips were fixed with 3.7 % paraformaldehyde for 10 min and washed twice with PBS and then permeabilised with 0.25 % Triton X-100 in PBS for 10 min at room temperature. Cells were blocked for 1 h with 5% goat serum in PBS-0.1% Triton. Primary antibody (diluted in blocking buffer) incubation was performed overnight at 4°C followed by secondary antibody conjugated to Alexa-488 or Alexa-594 incubation for 1 h at room temperature. Washes were performed with PBS-T (0.05% Tween20), between each antibody incubation. Nucleic acid staining was performed by 15 min incubation with Hoechst 33342 diluted 1:1,000 in PBS. For antibody dilutions, see the supplementary Table S2. For cell labelling using the recombinant EGFP-GRP1-PH protein, cells were permeabilised with 0.1% Triton X-100 in PBS and blocked in 3% fatty-acid free BSA and 0.05% Triton-X100 in PBS for 1 h at RT. This was followed by incubation with 40 μg/ml of the probe in 1% fatty-acid free BSA and 0.05% Triton-X100 in PBS for 2 h at RT. Images were acquired with a Leica DMI6000B fluorescence microscope using x40 or x100 objectives or Leica TCS SP5 confocal laser scanning microscope using a 63x/1.4 oil immersion lens. Images were processed with a Leica application suite V 4.0.

### SDS-PAGE and Western Immunoblotting

Proteins were resolved by SDS-PAGE and then transferred to nitrocellulose membranes. The membrane was then blocked with 7% milk in PBS-T (PBS pH 7.4, 0.05 % Tween-20) for 1 hour at room temperature before incubation with primary antibodies overnight at 4°C (for antibody dilutions see the supplementary Table S3). After washing with PBS-T, the membrane was incubated with HRP conjugated secondary antibodies for 1 hour at room temperature. The enhanced chemiluminescence (ThermoFisher Scientific) was added and the Chemidoc XRS+ imaging system from Bio-Rad was used for visualization.

### Recombinant protein expression and purification

The pGEX-4T1-EGFP-GRP1-PH plasmid was obtained from Julien Viaud (INSERM U.1048, Toulouse, France). The mutant K273A was generated by site-directed mutagenesis and validated by sequencing using ABI Prism BigDye Terminator version 3.1 cycle sequencing kit (Applied Biosystems) (see primer sequences in Supplementary Table S3). Following transformation into *E. coli* BL21-RIL DE3, the bacteria were grown at 37°C and further induced overnight at 18°C with 0.5 mM isopropyl-β-D-thiogalactopyranoside. Bacterial pellets were lysed in 50 mM Tris pH 7.5, 150 mM NaCl, 10 % de glycerol, 1% Triton-X100, 0.5 mg/ml lysozyme, 5 mM DTT and protease inhibitor cocktail for 30 min on ice. Following sonication and centrifugation, GST-EGFP-GRP1-PH was purified with glutathione-agarose 4B beads and analysed by SDS-PAGE and Coomassie staining. Expression and purification of GST-PLCδ1 was as described previously (46).

### Lipid overlay assay

Lipid overlay assays were performed according to Karlsson *et al* (25) using anti-PtdIns(3,4,5)*P*_3_ antibody followed by anti-mouse IgG-HRP secondary antibody or 0.5 to 1.5 μg/mL of recombinant GST (purified as described in (25)), GST-PARP1 (BPS Bioscience) followed by anti-GST-conjugated to HRP (1:30,000). Visualization was achieved with enhanced chemiluminescence. Binding of GST-EGFP-GRP1-PH was visualised by GFP fluorescence scanning using a Typhoon FLA 9000 scanner with an excitation of 473 nm and a BPB1 filter.

## Acknowledgments

We would like to thank Dr Bernd Thiede (University of Oslo, Norway) for validating the efficiency of the incorporation of heavy amino acids for SILAC labelling and Dr Diana C Turcu for excellent technical assistance. This project was funded by the University of Bergen, the Nansen Fond and the Meltzer foundation.

