## Supplementary tables S1-3 and Figure S1-3 for "Nuclear phosphatidylinositol 3,4,5-trisphosphate interactome uncovers an enrichment in nucleolar proteins"

#### Supplementary Tables S1-S3

#### Supplementary Figures S1-S3

### Supplementary Table S1. Nuclear PtdIns(3,4,5)*P*₃ interactome

Proteins pulled down specifically by PtdIns(3,4,5)$P_3$ identified with at least 2 peptides with heavy/light log2 ratio > 0.5 and common to the two replicate runs (179 proteins in total) are listed here. Validation of the identification in a second biological experiment is shown with a + in the last column.

ID names in bold are proteins previously experimentally reported to bind PtdIns(3,4,5)$P_3$.

proteins common to the T cell nucleolome list

proteins common with the nucleolar database (NoDB)

proteins common to the Lamont HeLa nucleolome list (2012)

GN: gene name

| Accession | Description | Coverage % | # Peptides | # PSMs | # Unique Peptides | # AAs | MW [kDa] | Abundance Ratio: Heavy / Light | log 2 ratio | (K/R-(Xn 3–7)-K-X-K/R-K/R) motif (bold) | Exp 2 |
|---|---|---|---|---|---|---|---|---|---|---|---|
| Q05639 | Elongation factor 1-alpha 2 GN=EEF1A2 | 16,2 | 7 | 18 | 1 | 463 | 50,438 | 100 | 6,6438 | | + |
| Q9BRX2 | Protein pelota homolog GN=PELO | 11,7 | 4 | 4 | 4 | 385 | 43,332 | **19,83** | **4,3096** | | |
| P29372 | DNA-3-methyladenine glycosylase GN=MPG | 19,1 | 4 | 4 | 4 | 298 | 32,848 | **18,812** | **4,2336** | | |
| P07900 | Heat shock protein HSP 90-alpha GN=HSP90AA1 | 5,3 | 3 | 4 | 3 | 732 | 84,607 | 13,62 | 3,7676 | 6:KEEEKEKEEKESEDKPEIEDVGSDEEEEK**KDGDKKKKKK**IKEKYIDQEELNKTKPIWTRN | |
| Q8NFW8 | N-acylneuraminate cytidylyltransferase GN=CMAS | 6,2 | 3 | 3 | 3 | 434 | 48,349 | **10,727** | **3,4232** | | |
| P08238 | Heat shock protein HSP 90-beta GN=HSP90AB1 | 11,0<br>4 | 6 | 7 | 6 | 724 | 83,212 | **10,343** | **3,3706** | 6:DKDDEEKPKIEDVGSDEEDDSG**KDKKKKTKK**IKEKYIDQEELNKTKPIWTRNPDDITQEE | |
| Q9UPQ0 | LIM and calponin homology domains-containing protein 1 GN=LIMCH1 | 3,7 | 3 | 3 | 3 | 1083 | 121,79 | **9,439** | **3,2386** | 12:SPLELKQDNGSIEINIKKPNSVPQELAATTEKTEPNSQED**KNDGGKSRK**GNIELASSEPQ | + |
| Q58FF8 | Putative heat shock protein HSP 90-beta 2 GN=HSP90AB2P | 5,8 | 2 | 2 | 2 | 381 | 44,321 | 8,559 | 3,0974 | 5:DDSG**KDKKKKTKK**IKEKYIDQEELNKTKPIWTRNTEDITQEEYGEFYKSLTNDWKDHLAV | |
| P35241 | Radixin GN=RDX | 6,7 | 4 | 11 | 1 | 583 | 68,521 | 5,822 | 2,5415 | 7:EVQQMKAQAREEKHQKQLE**RAQLENEKKKR**EIAEKEKERIEREKEELMERLKQIEEQTIK<br>8:AQKELEEQTRKALELDQERKRAKEEAE**RLEKERR**AAEEAKSAIAKQAADQMKNQEQLA<br>AE<br>9:LAEFTA**KIALLEEAKKKK**EEEATEWQHKAFAAQEDLEKTKEELKTVMSAPPPPPPPPVIP | |
| P35580 | Myosin-10 GN=MYH10 | 2,5 | 4 | 10 | 4 | 1976 | 228,86 | **5,623** | **2,4913** | 18:EEEGARQKLQLE**KVTAEAKIKK**MEEEILLLEDQNSKFIKEKKLMEDRIAECSSQLAEEEE<br>19:KAKNLAKIRNKQEVMISDLEERLKKEEKT**RQELEKAKRK**LDGETTDLQDQIAELQAQID<br>E<br>22:HATALEELSEQLEQAKRFKANLEKNKQGLETDNKELACEVKVLQQV**KAESEHKRKK**LD<br>AQ<br>27:EFERQNKQLRADMEDLMSSKDDVG**KNVHELEKSKR**ALEQQVEEMRTQLEELEDELQ<br>ATED | + |
| P35579 | Myosin-9 GN=MYH9 | 22,2 | 31 | 52 | 31 | 1960 | 226,39 | **5,071** | **2,3423** | 9:TYERMFRWLVLRINKALDKTKRQGASFIGILDIAGFEIFDLNSFEQLCINYTNEKLQQLF<br>18:KLQLEKVTTEAKLKKLEEEQIILEDQNCKLAKEKKLLEDRIAEFTTNLTEEEEKSKSLAK<br>19:LKNKHEAMITDLEERLRREEKQRQELEKTRRKLEGDSTDLSDQIAELQAQIAELKMQLA<br>K | + |

| | | | | | | | | | | | |
|---|---|---|---|---|---|---|---|---|---|---|---|
| | | | | | | | | | | 20:KEEELQAALARVEEEAAQKNMALKKIRELESQISELQEDLESERASRNKAEKQKRDLGEE<br>22:LAEQLEQTKRVKANLEKAKQTLENERGELANEVKVLLQGKGDSEHKRKKVEAQLQELQVK<br>27:QFRTEMEDLMSSKDDVGKSVHELEKSKRALEQQVEEMKTQLEELEDELQATEDAKLRLEV | |
| O94979 | Protein transport protein Sec31A GN=SEC31A | 2,5 | 3 | 3 | 3 | 1220 | 132,93 | **5,049** | **2,336** | | |
| P35749 | Myosin-11 GN=MYH11 | 2,8 | 4 | 4 | 4 | 1972 | 227,2 | **4,617** | **2,207** | 4:FCVVVNPYKHLPIYSEKIVDMYKGKKRHEMPPHIYAIADTAYRSMLQDREDQSILCTGES<br>18:EEEAARQKLQLEKVTAEAKIKKLEDEILVMDDQNNKLSKERKLLEERISDLTTNLAEEEE<br>19:KAKNLTKLKNKHESMISELEVRLKKEEKSRQELEKLKRKLEGDASDFHEQIADLQAQIAE<br>22:HAQAVEELTEQLEQFKRAKANLDKNKQTLEKENADLAGELRVLGQAKQEVEHKKKKLEAQ<br>27:ELERTNKMLKAEMEDLVSSKDDVGKNVHELEKSKRALETQMEEMKTQLEELEDELQATED<br>32:LHEMEGAVKSKFKSTIAALEAKIAQLEEQVEQEAREKQAATKSLKQKDKKLKEILLQVED | |
| P51116 | Fragile X mental retardation syndrome-related protein 2 GN=FXR2 | 9,2 | 5 | 5 | 4 | 673 | 74,178 | **4,28** | **2,0976** | | + |
| O00148 | ATP-dependent RNA helicase DDX39A GN=DDX39A | 21,3 | 8 | 12 | 3 | 427 | 49,098 | 4,105 | 2,0374 | 6:FMQDPMEVFVDDETKLTLHGLQQYYV**KLKDSEKNRK**LFDLLDVLEFNQVIIFVKSVQRCM | + |
| P15311 | Ezrin GN=EZR | 11,9 | 7 | 15 | 4 | 586 | 69,37 | **4,007** | **2,0025** | 7:EVQQMKAQAREEKHQKQLERQQLETEKKRRETVEREKEQMMREKEELMLRLQDYEEKTKK | |
| P67809 | Nuclease-sensitive element-binding protein 1 GN=YBX1 | 20,3 | 4 | 6 | 1 | 324 | 35,903 | 3,974 | 1,9906 | | + |
| Q7Z406 | Myosin-14 GN=MYH14 | 2,4 | 4 | 5 | 4 | 1995 | 227,73 | **3,962** | **1,9862** | 18:EKKRLQQHIQELEAHLEAEEGARQKLQLEKVTTEAKMKKFEEDLLLLEDQNSKLSKERKL<br>19:LEDRLAEFSSQAAEEEEKVKSLNKLRLKYEATIADMEDRLRKEEKGRQELEKLKRRLDGE<br>21:QEDLESERVARTKAEKQRRDLGEELEALRGELEDTLDSTNAQQELRSKREQEVTELKKTL | |
| Q92945 | Far upstream element-binding protein 2 GN=KHSRP | 7,9 | 4 | 4 | 4 | 711 | 73,07 | **3,831** | **1,9377** | | |
| Q13200 | 26S proteasome non-ATPase regulatory subunit 2 GN=PSMD2 | 5,7 | 4 | 4 | 4 | 908 | 100,14 | **3,706** | **1,8899** | 2:MEEGGRDKAPVQPQQSPAAAPGGTDE**KPSGKERR**DAGDKDKEQELSEEDKQLQDELEMLV<br>12:AGSGNVLKVQQLLHICSEHFDS**KEKEEDKDKK**EKKDKDKKEAPADMGAHQGVAVLGIALI | |
| P16989 | Y-box-binding protein 3 GN=YBX3 | 13,7 | 4 | 7 | 1 | 372 | 40,066 | 3,613 | 1,853197 | | + |
| P54886 | Delta-1-pyrroline-5-carboxylate synthase GN=ALDH18A1 | 9,2 | 6 | 6 | 6 | 795 | 87,248 | **3,509** | **1,8111** | | + |
| Q13838 | Spliceosome RNA helicase DDX39B GN=DDX39B | 21,3 | 8 | 13 | 3 | 428 | 48,96 | **3,301** | **1,7229** | 6:KFMQDPMEIFVDDETKLTLHGLQQYYV**KLKDNEKNRK**LFDLLDVLEFNQVVIFVKSVQRC | + |
| Q32P51 | Heterogeneous nuclear ribonucleoprotein A1-like 2 GN=HNRNPA1L2 | 26,2 | 9 | 22 | 7 | 320 | 34,204 | **3,272** | **1,7102** | | |
| P51114 | Fragile X mental retardation syndrome-related protein 1 GN=FXR1 | 14,6 | 7 | 8 | 6 | 621 | 69,678 | **3,229** | **1,6911** | 11:ISRAESQSRQRNLP**RETLAKNKK**EMAKDVIEEHGPSEKAINGPTSASGDDISKLQRTPGE | + |
| P42166 | Lamina-associated polypeptide 2, isoform alpha GN=TMPO | 3,4 | 2 | 3 | 2 | 694 | 75,446 | **3,096** | **1,6304** | | + |
| Q13112 | Chromatin assembly factor 1 subunit B GN=CHAF1B | 4,1 | 2 | 2 | 2 | 559 | 61,454 | **3,057** | **1,6121** | | |
| O75150 | E3 ubiquitin-protein ligase BRE1B GN=RNF40 | 4,6 | 4 | 4 | 4 | 1001 | 113,58 | **3,037** | **1,6026** | 2:MSGPGNKRAAGDGGSGPPEKKLSREEKTTTTLIEPIRLGGISSTEEMDLKVLQFKNKKLA<br>17:ESFNLKRAQEDISRLRRKLEKQRKVEVYADADEILQEEIKEYKARLTCPCCNTRKKDAVL | |
| P26599 | Polypyrimidine tract-binding protein 1 GN=PTBP1 | 31,6 | 9 | 36 | 9 | 531 | 57,186 | **2,991** | **1,5806** | 10:V**KGFKFFQKDRK**MALIQMGSVEEAVQALIDLHNHDLGENHHLRVSFSKSTI | |

| | | | | | | | | | | | |
|---|---|---|---|---|---|---|---|---|---|---|---|
| P53396 | ATP-citrate synthase GN=ACLY | 25,8 | 22 | 42 | 22 | 1101 | 120,76 | **2,985** | **1,5777** | | |
| P46060 | Ran GTPase-activating protein 1 GN=RANGAP1 | 8,5 | 4 | 4 | 4 | 587 | 63,502 | **2,952** | **1,5617** | | |
| Q08J23 | tRNA (cytosine(34)-C(5))-methyltransferase GN=NSUN2 | 2,2 | 2 | 2 | 2 | 767 | 86,416 | **2,905** | **1,5385** | | |
| Q96AE4 | Far upstream element-binding protein 1 GN=FUBP1 | 3,3 | 2 | 4 | 1 | 644 | 67,518 | 2,866 | 1,5191 | | |
| Q96I24 | Far upstream element-binding protein 3 GN=FUBP3 | 13,1 | 7 | 9 | 6 | 572 | 61,602 | **2,846** | **1,5089** | | |
| Q8WX93 | Palladin GN=PALLD | 3,8 | 5 | 6 | 5 | 1383 | 150,47 | **2,844** | **1,5079** | | + |
| P11586 | C-1-tetrahydrofolate synthase, cytoplasmic GN=MTHFD1 | 15,4 | 13 | 17 | 13 | 935 | 101,5 | **2,84** | **1,5059** | | + |
| **P46940** | **Ras GTPase-activating-like protein IQGAP1 GN=IQGAP1** | 17,0 | 27 | 41 | 27 | 1657 | 189,13 | **2,825** | **1,4983** | 25:ARTILLNTKRLIVDVIRFQPGETLTEILETPATSEQEAEHQRAMQRRAIRDA**KTPDKMKK**<br>27:AELVKLQQTYAALNSKATFYGEQVDYYKSYIKTCLDNLASKGKVS**KKPREMKGKK**SKKIS | + |
| P13010 | X-ray repair cross-complementing protein 5 GN=XRCC5 | 28,1 | 15 | 30 | 15 | 732 | 82,652 | **2,817** | **1,4942** | | |
| Q05193 | Dynamin-1 GN=DNM1 | 2,4 | 2 | 4 | 2 | 864 | 97,347 | **2,719** | **1,4431** | 11:YWFVLTAENLSWY**KDDEEKEKK**YMLSVDNLKLRDVEKGFMSSKHIFALFNTEQRNVYKDY | |
| P50570 | Dynamin-2 GN=DNM2 | 2,4 | 2 | 4 | 2 | 870 | 98,003 | **2,719** | **1,4431** | 11:YWFVLTAESLSWY**KDEEEKEKK**YMLPLDNLKIRDVEKGFMSNKHVFAIFNTEQRNVYKDL | + |
| Q9UQ16 | Dynamin-3 GN=DNM3 | 2,4 | 2 | 4 | 2 | 869 | 97,685 | **2,719** | **1,4431** | 11:KGGSKGYWFVLTAESLSWY**KDDEEKEKK**YMLPLDNLKVRDVEKSFMSSKHIFALFNTEQR | |
| O00571 | ATP-dependent RNA helicase DDX3X GN=DDX3X | 5,1 | 3 | 6 | 2 | 662 | 73,198 | **2,688** | **1,4265** | 5:SFSDVEMGEIIMGNIELTRYTRPTPVQ**KHAIPIIKEKR**DLMACAQTGSGKTAAFLLPILS | + |
| O15523 | ATP-dependent RNA helicase DDX3Y GN=DDX3Y | 5,1 | 3 | 6 | 2 | 660 | 73,108 | **2,688** | **1,4265** | 5:SDIDMGEIIMGNIELTRYTRPTPVQ**KHAIPIIKGKR**DLMACAQTGSGKTAAFLLPILSQI | |
| Q15019 | Septin-2 GN=SEPT2 | 29,6 | 7 | 11 | 7 | 361 | 41,461 | **2,647** | **1,4044** | | + |
| P60842 | Eukaryotic initiation factor 4A-I GN=EIF4A1 | 28,8 | 11 | 18 | 11 | 406 | 46,125 | **2,626** | **1,3929** | | + |
| Q99729 | Heterogeneous nuclear ribonucleoprotein A/B GN=HNRNPAB | 9,3 | 3 | 4 | 2 | 332 | 36,202 | **2,604** | **1,3807** | | |
| Q13283 | Ras GTPase-activating protein-binding protein 1 GN=G3BP1 | 24,5 | 8 | 16 | 8 | 466 | 52,132 | **2,589** | **1,3724** | | |
| O43719 | HIV Tat-specific factor 1 GN=HTATSF1 | 2,8 | 2 | 3 | 2 | 755 | 85,801 | **2,572** | **1,3629** | 5:KGDGLCCYLKRESVELALKLLDEDEIRGYKLHVEVAKFQL**KGEYDASKKKKK**CKDYKKKL | |
| P12814 | Alpha-actinin-1 GN=ACTN1 | 27,01 | 21 | 86 | 9 | 892 | 102,99 | **2,56** | **1,3561** | 5:ALIHRH**RPELIDYGKLRK**DDPLTNLNTAFDVAEKYLDIPKMLDAEDIVGTARPDEKAIMT | |
| Q7KZF4 | Staphylococcal nuclease domain-containing protein 1 GN=SND1 | 25,9 | 19 | 26 | 19 | 910 | 101,93 | **2,552** | **1,3516** | 7:IAVYTRGAEKL**RAAERFAKERR**LRIWRDYVAPTANLDQKDKQFVAKVMQVLNADAIVVKL | |
| Q00341 | Vigilin GN=HDLBP | 13,1 | 14 | 16 | 14 | 1268 | 141,37 | **2,548** | **1,3494** | 4:KDQGLSIMVSGKLDAVMKARKDIVARLQTQASATVAIPKEHHRFVIGKNGEKLQDLELKT<br>6:YNRLVGEIMQETGTRINIPPPSVNRTEIVFTGEKEQLAQAVARIKKIYEEKKKKTTTIAV<br>13:PAKLHNSLIGTKGRLIRSIMEECGGVHIHFPVEGSGSDTVVIRGPSSDVEKAKKQLLHLA<br>14:EEKQTKSFTVDIRAKPEYHKFLIGKGGGKIRKVRDSTGARVIFPAAEDKDQDLITIIGKE | |
| P62140 | Serine/threonine-protein phosphatase PP1-beta catalytic subunit GN=PPP1CB | 6,7 | 2 | 2 | 2 | 327 | 37,163 | 2,537 | 1,3431 | | + |
| P26038 | Moesin GN=MSN | 18,9 | 10 | 19 | 7 | 577 | 67,778 | **2,535** | **1,342** | 7:EVQQMKAQAREEKHQKQME**RAMLENEKKKR**EMAEKEKEKIEREKEELMERLKQIEEQTKK | + |
| P20042 | Eukaryotic translation initiation factor 2 subunit 2 GN=EIF2S2 | 12,6 | 3 | 4 | 3 | 333 | 38,364 | **2,505** | **1,3248** | 2:MSGDEMIFDPTMS**KKKKKKKK**PFMLDEEGDTQTEETQPSETKEVEPEPTEDKDLEADEED<br>3:TRKKDASDDLDDLNFFNQ**KKKKKK**TKKIFDIDEAEEGVKDLKIESDVQEPTEPEDDLDIM | + |

| | | | | | | | | | | | |
|---|---|---|---|---|---|---|---|---|---|---|---|
| P09874 | Poly [ADP-ribose] polymerase 1 GN=PARP1 | 26,8 | 22 | 37 | 22 | 1014 | 113,01 | **2,5** | **1,3219** | 3:GHSIRHPDVEVDGFSEL**RWDDQQKVKK**TAEAGGVTGKGQDGIGSKAEKTLGDFAAEYAKS | |
| Q2VIR3 | Putative eukaryotic translation initiation factor 2 subunit 3-like protein GN=EIF2S3L PE=5 SV=2 | 12,07 | 5 | 7 | 5 | 472 | 51,196 | **2,497** | **1,3202** | | |
| Q69YN4 | Protein virilizer homolog GN=KIAA1429 | 5,6 | 8 | 9 | 8 | 1812 | 201,9 | **2,474** | **1,3068** | | |
| Q92466 | DNA damage-binding protein 2 GN=DDB2 | 4,7 | 2 | 2 | 2 | 427 | 47,833 | **2,474** | **1,3068** | | |
| Q14240 | Eukaryotic initiation factor 4A-II GN=EIF4A2 | 16,5 | 6 | 11 | 6 | 407 | 46,373 | **2,459** | **1,2981** | | |
| Q8NE71 | ATP-binding cassette sub-family F member 1 GN=ABCF1 | 5,8 | 4 | 5 | 4 | 845 | 95,866 | **2,426** | **1,2786** | 2:MPKAPKQQPPEPEWIGDGESTSPSDKVVKKGKKDKKIKKTFFEELAVEDKQAGEEEKVLK<br>3:EKEQQQQQQQQQKKKRDTRKGRRKKDVDDDGEEKELMERLKKLSVPTSDEEDEVPAPKP<br>5:EKSKGKAKPQNKFAALDNEEEDKEEEIIKEKEPPKQGKEKAKKAEQGSEEEGEGEEEEEE<br>6:GGESKADDPYAHLSKKEKKKLKKQMEYERQVASLKAANAAENDFSVSQAEMSSRQAMLEN<br>11:QQKQKELLKQYEKQEKKLKELKAGGKSTKQAEKQTKEALTRKQQKCRRKNQDEESQEAPE | |
| Q9NSD9 | Phenylalanine--tRNA ligase beta subunit GN=FARSB | 15,9 | 9 | 13 | 9 | 589 | 66,074 | **2,415** | **1,272** | | |
| Q52LJ0 | Protein FAM98B GN=FAM98B | 8,2 | 2 | 2 | 2 | 330 | 37,167 | **2,408** | **1,2678** | | + |
| O75475 | PC4 and SFRS1-interacting protein GN=PSIP1 | 3,2 | 2 | 2 | 2 | 530 | 60,067 | **2,407** | **1,2672** | | |
| Q13263 | Transcription intermediary factor 1-beta GN=TRIM28 | 12,1 | 8 | 9 | 8 | 835 | 88,493 | **2,383** | **1,2528** | | |
| Q8N684 | Cleavage and polyadenylation specificity factor subunit 7 GN=CPSF7 | 16,1 | 5 | 6 | 5 | 471 | 52,018 | **2,377** | **1,2491** | | |
| P52272 | Heterogeneous nuclear ribonucleoprotein M GN=HNRNPM | 25,9 | 14 | 21 | 14 | 730 | 77,464 | **2,37** | **1,2449** | | + |
| P41091 | Eukaryotic translation initiation factor 2 subunit 3 GN=EIF2S3 | 15,04 | 6 | 8 | 6 | 472 | 51,077 | **2,367** | **1,2431** | | |
| O43707 | Alpha-actinin-4 GN=ACTN4 | 65,7 | 46 | 175 | 34 | 911 | 104,79 | **2,366** | **1,2425** | 5:KNVNVQNFHISWKDGLAFNALIHRHRPELIEYDKLRKDDPVTNLNNAFEVAEKYLDIPKM | |
| P0DMV9 | Heat shock 70 kDa protein 1B GN=HSPA1B | 17,6 | 9 | 19 | 7 | 641 | 70,009 | **2,355** | **1,2357** | | |
| Q14103 | Heterogeneous nuclear ribonucleoprotein D0 GN=HNRNPD | 12,7 | 5 | 8 | 4 | 355 | 38,41 | **2,345** | **1,2296** | | |
| P23284 | Peptidyl-prolyl cis-trans isomerase B GN=PPIB | 31,5 | 7 | 10 | 7 | 216 | 23,728 | **2,342** | **1,2277** | | + |
| Q9P258 | Protein RCC2 GN=RCC2 | 30,6 | 14 | 18 | 14 | 522 | 56,049 | **2,305** | **1,2048** | | + |
| Q09028 | Histone-binding protein RBBP4 GN=RBBP4 | 15,5 | 6 | 7 | 6 | 425 | 47,626 | **2,305** | **1,2048** | | |
| P09651 | Heterogeneous nuclear ribonucleoprotein A1 GN=HNRNPA1 | 26,9 | 10 | 24 | 8 | 372 | 38,723 | **2,286** | **1,1928** | | + |
| P46063 | ATP-dependent DNA helicase Q1 GN=RECQL | 10,01 | 5 | 7 | 5 | 649 | 73,41 | **2,271** | **1,1833** | 12:ESSQTCHSEQGDKKMEEKNSGNFQKKAANMLQQSGS**KNTGAKKRK**IDDA | |
| P48741 | Putative heat shock 70 kDa protein 7 GN=HSPA7 PE=5 SV=2 | 13,6 | 4 | 10 | 2 | 367 | 40,22 | **2,269** | **1,1821** | | |
| P33991 | DNA replication licensing factor MCM4 GN=MCM4 | 6,02 | 5 | 6 | 5 | 863 | 96,498 | **2,264** | **1,1789** | | |
| P05198 | Eukaryotic translation initiation factor 2 subunit 1 GN=EIF2S1 | 23,5 | 7 | 10 | 7 | 315 | 36,089 | **2,249** | **1,1693** | 4:YTKDEQLESLFQRTAWVFDDKYKRPGYGAYDAFKHAVSDPSILDSLDLNEDEREVLINNI | |
| Q6S8J3 | POTE ankyrin domain family member E GN=POTEE | 3,7 | 3 | 7 | 3 | 1075 | 121,29 | **2,243** | **1,1654** | | |
| A5A3E0 | POTE ankyrin domain family member F GN=POTEF | 3,7 | 3 | 7 | 3 | 1075 | 121,37 | **2,243** | **1,1654** | | |

| | | | | | | | | | | | |
|---|---|---|---|---|---|---|---|---|---|---|---|
| P60709 | Actin, cytoplasmic 1 GN=ACTB | 34,9 | 9 | 18 | 9 | 375 | 41,71 | **2,241** | **1,1641** | | + |
| P63261 | Actin, cytoplasmic 2 GN=ACTG1 | 34,9 | 9 | 18 | 9 | 375 | 41,766 | **2,241** | **1,1641** | | |
| P78347 | General transcription factor II-I GN=GTF2I | 9,02 | 8 | 12 | 8 | 998 | 112,35 | **2,236** | **1,1609** | | |
| Q14157 | Ubiquitin-associated protein 2-like GN=UBAP2L | 2,1 | 2 | 2 | 2 | 1087 | 114,47 | **2,216** | **1,148** | | + |
| P12956 | X-ray repair cross-complementing protein 6 GN=XRCC6 | 38,1 | 18 | 40 | 18 | 609 | 69,799 | **2,215** | **1,1473** | | |
| P78527 | DNA-dependent protein kinase catalytic subunit GN=PRKDC | 14,00 | 50 | 74 | 50 | 4128 | 468,79 | **2,212** | **1,1454** | 21:**KHVSLNKAKKRR**LPRGFPPSASLCLLDLVKWLLAHCGRPQTECRHKSIELFYKFVPLLPG | + |
| P62826 | GTP-binding nuclear protein Ran GN=RAN | 14,8 | 3 | 3 | 3 | 216 | 24,408 | **2,205** | **1,1408** | 4:GN**KVDIKDRK**VKAKSIVFHRKKNLQYYDISAKSNYNFEKPFLWLARKLIGDPNLEFVAMP | |
| O43684 | Mitotic checkpoint protein BUB3 GN=BUB3 | 19,5 | 5 | 13 | 5 | 328 | 37,131 | **2,164** | **1,1137** | | |
| Q05048 | Cleavage stimulation factor subunit 1 GN=CSTF1 | 14,8 | 5 | 6 | 5 | 431 | 48,327 | **2,146** | **1,1017** | | + |
| Q15393 | Splicing factor 3B subunit 3 GN=SF3B3 | 5,1 | 5 | 5 | 5 | 1217 | 135,49 | **2,139** | **1,0969** | | + |
| P54652 | Heat shock-related 70 kDa protein 2 GN=HSPA2 | 11,9 | 7 | 15 | 4 | 639 | 69,978 | **2,136** | **1,0949** | | |
| P49916 | DNA ligase 3 GN=LIG3 | 6,3 | 6 | 6 | 6 | 1009 | 112,84 | **2,119** | **1,0834** | 13:DVKGTYEPG**KRHWLKVKK**DYLNEGAMADTADLVVLGAFYGQGSKGGMMSIFLMGCYDPGS | |
| Q16531 | DNA damage-binding protein 1 GN=DDB1 | 4,5 | 5 | 6 | 5 | 1140 | 126,89 | **2,101** | **1,0711** | | |
| Q15428 | Splicing factor 3A subunit 2 GN=SF3A2 | 5,6 | 2 | 3 | 2 | 464 | 49,224 | **2,094** | **1,0663** | | |
| Q9BYX7 | Putative beta-actin-like protein 3 GN=POTEKP PE=5 SV=1 | 7,7 | 2 | 6 | 2 | 375 | 41,989 | **2,092** | **1,0649** | | |
| P26358 | DNA (cytosine-5)-methyltransferase 1 GN=DNMT1 | 2,9 | 5 | 5 | 5 | 1616 | 183,05 | **2,079** | **1,0559** | 5:QEESERAKSDESIKEEDKDQDEKRRRVTSRERVARPLPAEEPERAKSGTRTEKEEERDEK<br>7:EKEPEKVNPQISDEKDEDEKEEKRRKTTPKEPTEKKMARAKTVMNSKTHPPKCIQCGQYL<br>12:RQTIRHSTREKDRGPTKATTTKLVYQIFDTFFAEQIEKDDREDKENAFKRRRCGVCEVCQ | |
| P29692 | Elongation factor 1-delta GN=EEF1D | 26,7 | 5 | 7 | 5 | 281 | 31,103 | **2,078** | **1,0552** | | + |
| Q8N163 | Cell cycle and apoptosis regulator protein 2 GN=CCAR2 | 3,5 | 3 | 4 | 3 | 923 | 102,84 | **2,078** | **1,0552** | 5:FPARGPHGRLDQG**RSDDYDSKKRK**QRAGGEPWGAKKPRHDLPPYRVHLTPYTVDSPICDF | |
| Q8NBJ5 | Procollagen galactosyltransferase 1 GN=COLGALT1 | 8,8 | 5 | 8 | 5 | 622 | 71,59 | **2,069** | **1,0489** | | + |
| Q1KMD3 | Heterogeneous nuclear ribonucleoprotein U-like protein 2 GN=HNRNPUL2 | 19,8 | 11 | 20 | 11 | 747 | 85,052 | **2,036** | **1,0257** | | + |
| O43670 | BUB3-interacting and GLEBS motif-containing protein ZNF207 GN=ZNF207 | 5,02 | 2 | 3 | 2 | 478 | 50,717 | **2,025** | **1,0179** | | |
| Q01780 | Exosome component 10 GN=EXOSC10 | 3,4 | 2 | 2 | 2 | 885 | 100,77 | **2,009** | **1,0065** | 5:IRPQLKFREKIDNSNTPFLPKIFIKPNAQ**KPLPQALSKERR**ERPQDRPEDLDVPPALADF | |
| P68104 | Elongation factor 1-alpha 1 GN=EEF1A1 | 27,5 | 9 | 40 | 3 | 462 | 50,109 | **2,007** | **1,005** | | |
| Q5VTE0 | Putative elongation factor 1-alpha-like 3 GN=EEF1A1P5 | 27,5 | 9 | 40 | 3 | 462 | 50,153 | **2,007** | **1,005** | | |
| P68032 | Actin, alpha cardiac muscle 1 GN=ACTC1 | 10,1 | 3 | 5 | 3 | 377 | 41,992 | **2,003** | **1,0022** | | + |
| P68133 | Actin, alpha skeletal muscle GN=ACTA1 | 10,1 | 3 | 5 | 3 | 377 | 42,024 | **2,003** | **1,0022** | | |
| P62736 | Actin, aortic smooth muscle GN=ACTA2 | 10,1 | 3 | 5 | 3 | 377 | 41,982 | **2,003** | **1,0022** | | |
| P63267 | Actin, gamma-enteric smooth muscle GN=ACTG2 | 10,1 | 3 | 5 | 3 | 376 | 41,85 | **2,003** | **1,0022** | | |
| P11142 | Heat shock cognate 71 kDa protein GN=HSPA8 | 30,6 | 17 | 33 | 14 | 646 | 70,854 | **1,999** | **0,9993** | | |
| Q9UN86 | Ras GTPase-activating protein-binding protein 2 GN=G3BP2 | 8,1 | 3 | 6 | 3 | 482 | 54,088 | **1,999** | **0,9993** | | + |

| | | | | | | | | | | | |
|---|---|---|---|---|---|---|---|---|---|---|---|
| Q9BR76 | Coronin-1B GN=CORO1B | 12,2 | 6 | 7 | 6 | 489 | 54,2 | **1,988** | **0,9913** | | |
| Q9Y3I0 | tRNA-splicing ligase RtcB homolog GN=RTCB | 23,2 | 10 | 15 | 10 | 505 | 55,175 | **1,971** | **0,9789** | 9:ETFGTTCHGAG**RALSRAKSRR**NLDFQDVLDKLADMGIAIRVASPKLVMEEAPESYKNVTD | + |
| P62136 | Serine/threonine-protein phosphatase PP1-alpha catalytic subunit GN=PPP1CA | 13,9 | 4 | 4 | 4 | 330 | 37,488 | 1,939 | 0,9553 | 7:KNKGKYGQFSGLNPGGRPITPP**RNSAKAKK** | |
| P50552 | Vasodilator-stimulated phosphoprotein GN=VASP | 12,4 | 4 | 7 | 4 | 380 | 39,805 | **1,927** | **0,9464** | | |
| Q16181 | Septin-7 GN=SEPT7 | 12,1 | 4 | 8 | 4 | 437 | 50,648 | **1,915** | **0,9373** | 7:TNNVHYENYRSRKLAAVTYNGVDNNKNKGQLTKSPLAQMEEE**RREHVAKMKK**MEMEMEQV<br>9:QNSS**RTLEKNKKK**GKIF | |
| Q14444 | Caprin-1 GN=CAPRIN1 | 9,3 | 6 | 9 | 6 | 709 | 78,318 | **1,911** | **0,9343** | | |
| Q9NVA2 | Septin-11 GN=SEPT11 | 10,2 | 4 | 5 | 4 | 429 | 49,367 | **1,911** | **0,9343** | | |
| Q15007 | Pre-mRNA-splicing regulator WTAP GN=WTAP | 6,06 | 2 | 2 | 2 | 396 | 44,217 | **1,907** | **0,9313** | | |
| P14866 | Heterogeneous nuclear ribonucleoprotein L GN=HNRNPL | 20,03 | 7 | 10 | 7 | 589 | 64,092 | **1,895** | **0,9222** | 2:MS**RRLLPRAEKRRRR**LEQRQQPDEQRRRSGAMVKMAAAGGGGGGGRYYGGGSEGGRAPKR | + |
| Q12874 | Splicing factor 3A subunit 3 GN=SF3A3 | 16,6 | 6 | 10 | 6 | 501 | 58,812 | **1,873** | **0,9054** | | |
| Q12996 | Cleavage stimulation factor subunit 3 GN=CSTF3 | 6,4 | 4 | 7 | 4 | 717 | 82,869 | **1,855** | **0,8914** | | + |
| Q15459 | Splicing factor 3A subunit 1 GN=SF3A1 | 20,8 | 14 | 20 | 14 | 793 | 88,831 | **1,854** | **0,8906** | 5:FLTQLMQKEQRNYQFDFLRPQHSLFNYFTKLVEQYTKILIPP**KGLFSKLKK**EAENPREVL | |
| P42285 | Superkiller viralicidic activity 2-like 2 GN=SKIV2L2 | 4,6 | 4 | 5 | 4 | 1042 | 117,73 | **1,85** | **0,8875** | 19:N**KFAEGITKIKR**DIVFAASLYL | |
| Q92499 | ATP-dependent RNA helicase DDX1 GN=DDX1 | 25,5 | 16 | 24 | 16 | 740 | 82,38 | **1,846** | **0,8844** | | + |
| Q14676 | Mediator of DNA damage checkpoint protein 1 GN=MDC1 | 0,9 | 2 | 2 | 2 | 2089 | 226,53 | **1,832** | **0,8734** | 6:ATVEAKQSEAEVVTEIQLEKDQPLV**KERDNDTKVKR**GAGNGVVPAGVILERSQPPGEDSD | |
| Q14315 | Filamin-C GN=FLNC | 4,8 | 11 | 26 | 6 | 2725 | 290,84 | **1,829** | **0,8711** | | + |
| Q01844 | RNA-binding protein EWS GN=EWSR1 | 7,3 | 3 | 5 | 3 | 656 | 68,436 | **1,806** | **0,8528** | | |
| Q00610 | Clathrin heavy chain 1 GN=CLTC | 9,3 | 12 | 21 | 12 | 1675 | 191,49 | **1,794** | **0,8432** | | + |
| P11021 | 78 kDa glucose-regulated protein GN=HSPA5 | 20,9 | 10 | 19 | 8 | 654 | 72,288 | **1,787** | **0,8375** | 6:VFEVVATNGDTHLGGEDFDQRVMEHFIKLYKKKTGKDV**RKDNRAVQKLRR**EVEKAKRALS | + |
| O00303 | Eukaryotic translation initiation factor 3 subunit F GN=EIF3F | 15,7 | 4 | 6 | 4 | 357 | 37,54 | **1,785** | **0,8359** | | + |
| P38646 | Stress-70 protein, mitochondrial GN=HSPA9 | 5,4 | 3 | 4 | 3 | 679 | 73,635 | **1,777** | **0,8294** | | |
| O14979 | Heterogeneous nuclear ribonucleoprotein D-like GN=HNRNPDL | 7,8 | 3 | 4 | 2 | 420 | 46,409 | 1,775 | 0,8278 | | + |
| Q14683 | Structural maintenance of chromosomes protein 1A GN=SMC1A | 6,1 | 7 | 10 | 7 | 1233 | 143,14 | **1,772** | **0,8254** | 6:ELASKNKEIEKDKKRMDKVEDELKEKKKELGKMMREQQQIEKEIKEKDSELNQKRPQYIK<br>7:AKENTSHKIKKLEAAKKSLQNAQKHYKKRKGDMDELEKEMLSVEKARQEFEERMEEESQS<br>13:AKARRWDEKAVDKLKEKKERLTEELKEQMKAKRKEAELRQVQSQAHGLQMRLKYSQSDLE<br>15:CREIGVRNIREFEEEKVKRQNEIAKKRLEFENQKTRLGIQLDFEKNQLKEDQDKVHMWEQ<br>16:TVKKDENEIEKLKKEEQRHMKIIDETMAQLQDLKNQHLAKKSEVNDKNHEMEEIRKKLGG | + |

| | | | | | | | | | | | |
|---|---|---|---|---|---|---|---|---|---|---|---|
| Q9UMS4 | Pre-mRNA-processing factor 19 GN=PRPF19 | 18,6 | 6 | 11 | 6 | 504 | 55,146 | **1,766** | **0,8205** | | + |
| Q15717 | ELAV-like protein 1 GN=ELAVL1 | 15,9 | 5 | 6 | 5 | 326 | 36,069 | **1,762** | **0,8172** | | |
| P25205 | DNA replication licensing factor MCM3 GN=MCM3 | 2,1 | 2 | 3 | 2 | 808 | 90,924 | **1,757** | **0,8131** | 13:EKKRKKRSEDESETEDEEE**KSQEDQEQKRKRRK**TRQPDAKDGDSYDPYDFSDTEEEMPQV | |
| P43243 | Matrin-3 GN=MATR3 | 6,8 | 4 | 8 | 4 | 847 | 94,565 | **1,743** | **0,8016** | 9:PFGVISNHLILNKINEAFIEMATTEDAQAAVDYYTTTPALVFGKPVRVHLSQ**KYKRIKK**P<br>11:TREDAMAMVDHCLKKALWFQGRCVKVDLSEKYKKLVLRIPNRGIDLL**KKDKSRKR**SYSPD | |
| Q00839 | Heterogeneous nuclear ribonucleoprotein U GN=HNRNPU | 19,9 | 14 | 44 | 14 | 825 | 90,528 | **1,742** | **0,8007** | 2:MSSSPVNVKKL**KVSELKEELKKRR**LSDKGLKAELMERLQAALDDEEAGGRPAMEPGNGSL | + |
| Q9Y285 | Phenylalanine--tRNA ligase alpha subunit GN=FARSA | 16,7 | 7 | 10 | 7 | 508 | 57,528 | **1,74** | **0,7991** | | |
| **O15294** | **UDP-N-acetylglucosamine--peptide N-acetylglucosaminyltransferase 110 kDa subunit GN=OGT** | 6,2 | 6 | 6 | 6 | 1046 | 116,85 | **1,725** | **0,7866** | | |
| P22626 | Heterogeneous nuclear ribonucleoproteins A2/B1 GN=HNRNPA2B1 | 35,4 | 11 | 25 | 9 | 353 | 37,407 | **1,723** | **0,7849** | | + |
| Q10567 | AP-1 complex subunit beta-1 GN=AP1B1 | 3,9 | 3 | 4 | 3 | 949 | 104,57 | **1,719** | **0,7816** | 2:MTDSKYFTTTKKGEIFEL**KAELNSDKKEKKK**EAVKKVIASMTVGKDVSALFPDVVNCMQT | |
| Q86YP4 | Transcriptional repressor p66-alpha GN=GATAD2A | 7,4 | 3 | 4 | 2 | 633 | 68,021 | **1,716** | **0,779** | 10:TAPAQAKAEPTAAPHPVL**KQVIKPRRK**LAFRSGEARDWSNGAVLQASSQLSRGSATTPRG | + |
| O75534 | Cold shock domain-containing protein E1 GN=CSDE1 | 3,7 | 2 | 2 | 2 | 798 | 88,829 | **1,708** | **0,7723** | | + |
| Q05682 | Caldesmon GN=CALD1 | 10,3 | 6 | 6 | 6 | 793 | 93,175 | **1,705** | **0,7698** | 5:EENKKED**KEKEEEEEEKPKR**GSIGENQVEVMVEEKTTESQEETVVMSLKNGQISSEEPKQ<br>9:KKAQEDKLQTAVLKKQGEEKGTKVQAKREKLQEDKPTFK**KEEIKDEKIKK**DKEPKEEVKS | + |
| P46821 | Microtubule-associated protein 1B GN=MAP1B | 4,5 | 9 | 9 | 9 | 2468 | 270,47 | **1,698** | **0,7638** | 8:VPENLKNPEPNIKMKRSIEEACFTLQYLNKLSMKPEPLFRSVGNTIDPVILFQKMGVGKL<br>12:SVTEKEVPSKEEPSPVKAEVAEKQATDVKPKAAKEKTVKKETKVKPEDKKEEKEKPKKEV<br>13:AKKEDKTPIKKEEKPKKEEVKKEVKKEIKKEEKKEPKKEVKKETPPKEVKKEVKKEEKKE<br>39:DPEALAIEQNLGKALKKDLKEKTKTKKPGTKTKSSSPVKKSDGKSKPLAASPKPAGLKES | + |
| P49736 | DNA replication licensing factor MCM2 GN=MCM2 | 8,2 | 6 | 9 | 6 | 904 | 101,83 | **1,68** | **0,7485** | | + |
| P55072 | Transitional endoplasmic reticulum ATPase GN=VCP | 8,3 | 6 | 6 | 6 | 806 | 89,266 | **1,672** | **0,7416** | | |
| Q02809 | Procollagen-lysine,2-oxoglutarate 5-dioxygenase 1 GN=PLOD1 | 8,7 | 5 | 7 | 5 | 727 | 83,497 | **1,651** | **0,7233** | | + |
| Q7L014 | Probable ATP-dependent RNA helicase DDX46 GN=DDX46 | 2,03 | 2 | 2 | 2 | 1031 | 117,29 | **1,648** | **0,7207** | 3:RSRDRKRLRRSRSRERDRSRERRRSRSRDRRRSRSRSRGRRSRSSSPGNKSKKTENRSRS<br>4:KE**KTDGGESSKEKK**KDKDDKEDEKEKDAGNFDQNKLEEEMRKRKERVEKWREEQRKKAME | |
| **P06748** | **Nucleophosmin GN=NPM1** | 18,02 | 3 | 15 | 3 | 294 | 32,555 | **1,639** | **0,7128** | | + |
| O14776 | Transcription elongation regulator 1 GN=TCERG1 | 1,8 | 2 | 2 | 2 | 1098 | 123,82 | **1,634** | **0,7084** | 13:ARMKQFKDMLLERGVSAFSTWEKELHKIVFDP**RYLLLNPKERK**QVFDQYVKTRAEEERRE | |
| Q16630 | Cleavage and polyadenylation specificity factor subunit 6 GN=CPSF6 | 17,06 | 6 | 9 | 6 | 551 | 59,173 | **1,632** | **0,7066** | 10:ESKSYGSGSRRERSRE**RDHSRSREKSRR**HKSRSRDRHDDYYRERSRERERHRDRDRDRDR | + |
| P13639 | Elongation factor 2 GN=EEF2 | 9,9 | 9 | 11 | 9 | 858 | 95,277 | **1,631** | **0,7058** | | |
| P51610 | Host cell factor 1 GN=HCFC1 | 4,3 | 7 | 10 | 7 | 2035 | 208,6 | **1,626** | **0,7013** | | |
| Q9UHD8 | Septin-9 GN=SEPT9 | 23,5 | 11 | 18 | 11 | 586 | 65,361 | **1,616** | **0,6924** | | |
| P60228 | Eukaryotic translation initiation factor 3 subunit E GN=EIF3E | 9,2 | 4 | 5 | 4 | 445 | 52,187 | **1,599** | **0,6772** | | + |
| Q13347 | Eukaryotic translation initiation factor 3 subunit I GN=EIF3I | 19,1 | 6 | 7 | 6 | 325 | 36,479 | **1,592** | **0,6708** | | + |
| P02545 | Prelamin-A/C GN=LMNA | 44,7 | 28 | 54 | 28 | 664 | 74,095 | **1,585** | **0,6645** | | + |

| | | | | | | | | | | | |
|---|---|---|---|---|---|---|---|---|---|---|---|
| O60934 | Nibrin GN=NBN | 3,2 | 2 | 2 | 2 | 754 | 84,906 | **1,575** | **0,6554** | 13:SLVIKNSTSRNPSGINDDYGQLKNFKKFKKVTYPGAGKLPHIIGGSDLIAHHARKNTELE<br>14:EWLRQEMEVQNQHAKEESLADDLFRYNPYLKRRR | + |
| Q92841 | Probable ATP-dependent RNA helicase DDX17 GN=DDX17 | 23,6 | 15 | 24 | 9 | 729 | 80,222 | **1,569** | **0,6498** | | + |
| Q9Y262 | Eukaryotic translation initiation factor 3 subunit L GN=EIF3L | 14,5 | 8 | 9 | 8 | 564 | 66,684 | **1,568** | **0,6489** | | + |
| P26641 | Elongation factor 1-gamma GN=EEF1G | 24,5 | 10 | 28 | 10 | 437 | 50,087 | **1,565** | **0,6462** | | + |
| **Q86V81** | **THO complex subunit 4 GN=ALYREF** | 24,5 | 3 | 6 | 3 | 257 | 26,872 | **1,56** | **0,6415** | | + |
| Q12905 | Interleukin enhancer-binding factor 2 GN=ILF2 | 11,5 | 4 | 12 | 4 | 390 | 43,035 | **1,552** | **0,6341** | | + |
| P33992 | DNA replication licensing factor MCM5 GN=MCM5 | 9,4 | 5 | 8 | 5 | 734 | 82,233 | **1,538** | **0,6211** | | |
| Q14152 | Eukaryotic translation initiation factor 3 subunit A GN=EIF3A | 5,7 | 7 | 9 | 7 | 1382 | 166,47 | **1,537** | **0,6201** | 13:LEELDPDFIMAKQVEQLEKEKKELQERLKNQE**KKIDYFERAKR**LEEIPLIKSAYEEQRIK | + |
| P61978 | Heterogeneous nuclear ribonucleoprotein K GN=HNRNPK | 35,4 | 13 | 36 | 13 | 463 | 50,944 | **1,532** | **0,6154** | | + |
| Q99613 | Eukaryotic translation initiation factor 3 subunit C GN=EIF3C | 9,3 | 8 | 12 | 8 | 913 | 105,28 | **1,528** | **0,6116** | 2:MSRFFTTGSDSESESSLSGEELVTKPVGGNYGKQPLLLSEDEEDTKRVVRSAKDKRFEEL<br>4:WEDKEGKKKMNKNNAKALSTLRQKIRKYNRDFESHITSYKQNPEQSADEDAEKNEEDS<br>EG<br>6:DSEEEEGKQTALASRFLKKAPTTDEDKKAAEKKREDKAKKKHDRKSKRLDEEEEDNEGGE | + |
| B5ME19 | Eukaryotic translation initiation factor 3 subunit C-like protein GN=EIF3CL | 9,3 | 8 | 12 | 8 | 914 | 105,41 | **1,528** | **0,6116** | 2:MSRFFTTGSDSESESSLSGEELVTKPVGGNYGKQPLLLSEDEEDTKRVVRSAKDKRFEEL<br>4:WEDKEGKKKMNKNNAKALSTLRQKIRKYNRDFESHITSYKQNPEQSADEDAEKNEEDS<br>EG<br>6:DSEEEEGKQTALASRFLKKAPTTDEDKKAAEKKREDKAKKKHDRKSKRLDEEEEEDNEGG | |
| Q96KR1 | Zinc finger RNA-binding protein GN=ZFR | 4,0 | 3 | 3 | 3 | 1074 | 116,94 | **1,52** | **0,6041** | 7:AAWTGTTFTKKAPFQNKQLKPKQPPKPPQIHYCDVCKISCAGPQTYKEHLEGQKHKKKE<br>A<br>12:LKGRRHRLQYKKKVNPDLQVEVKPSIRARKIQEEKMRKQMQKEEYWRRREEEERWR<br>MEMR<br>19:LAFRQIHKVLGMDPLPQMSQRFNIHNNRKRRRDSDGVDGFEAEGKKDKKDYDNF | + |
| P33993 | DNA replication licensing factor MCM7 GN=MCM7 | 15,8 | 9 | 10 | 9 | 719 | 81,257 | **1,516** | **0,6003** | 2:MAL**KDYALEKEKVKK**FLQEFYQDDELGKKQFKYGNQLVRLAHREQVALYVDLDDVAED<br>DP | + |
| P35637 | RNA-binding protein FUS GN=FUS | 18,1 | 6 | 20 | 6 | 526 | 53,394 | **1,513** | **0,5974** | | |
| O60506 | Heterogeneous nuclear ribonucleoprotein Q GN=SYNCRIP | 39,8 | 19 | 45 | 14 | 623 | 69,56 | **1,504** | **0,5888** | 8:FG**KLERVKK**LKDYAFIHFDERDGAVKAMEEMNGKDLEGENIEIVFAKPPDQKRKERKAQ<br>R | |
| P51659 | Peroxisomal multifunctional enzyme type 2 GN=HSD17B4 | 11,7 | 7 | 8 | 7 | 736 | 79,636 | **1,504** | **0,5888** | | |
| Q92804 | TATA-binding protein-associated factor 2N GN=TAF15 | 3,9 | 2 | 4 | 2 | 592 | 61,793 | **1,487** | **0,5724** | | |
| O96019 | Actin-like protein 6A GN=ACTL6A | 21,2 | 7 | 7 | 7 | 429 | 47,43 | **1,467** | **0,5529** | | + |
| Q13435 | Splicing factor 3B subunit 2 GN=SF3B2 | 6,4 | 5 | 6 | 5 | 895 | 100,17 | **1,465** | **0,5509** | 7:RSSLGQSASETEEDTVSVS**KKEKNRKRRNRKKKKK**PQRVRGVSSESSGDREKDSTRSRG<br>S<br>9:KKGFEEEHKDSDDDSSDDEQEKKPEAPKLSKKKLRRMNRFTVAELKQLVARPDVVEMH<br>DV<br>16:TQKYEEHVREQQAQVEKEDFSDMVAEHAAKQKQKKRKAQPQDSRGGSKKYKEFKF | + |
| Q86XP3 | ATP-dependent RNA helicase DDX42 GN=DDX42 | 8,3 | 6 | 8 | 6 | 938 | 102,91 | **1,463** | **0,5489** | 4:AEVEDQAARDMKRLEEKDKERKNVKGIRDDIEEEDDQEAYFRYMAENPTAGVVQEEEE<br>DN<br>12:LLTPKDSNFAGDLVRNLEGANQHVSKELLDLAMQNAWFRKSRFKGGKGKKLNIGGG<br>GLGY | + |

#### **Supplementary Table S2-** List of antibodies used in this study

IMF: immunofluorescence staining, LOA: lipid overlay assay and WB: Western immunoblotting.

| Antibodies | Reference Number | Company name | Dilution |
|---|---|---|---|
| **GST-HRP** | ab3416 | abcam | **LOA:** 1:10,000 |
| **Nucleolin** | 12247<br>14574S | Cell signaling Technology | **IMF:** 1:100<br>**IMF:** 1:100 |
| **PARP1** | 9542S | Cell signaling Technology | **IMF:** 1:50 |
| **PtdIns(3,4,5)$P_3$** | Z-P345b | Echelon | **IMF:** 1:400 |
| **UBF** | sc-9131 | Santacruz | **IMF:** 1:50 |
| **Goat anti-Mouse IgG Alexa Fluor 594** | A-11005 | Thermo Fisher Scientific | **IMF**: 1:200 |
| **Goat anti-Rabbit Alexa Fluor 594** | A-11012 | Thermo Fisher Scientific | **IMF**: 1:200 |
| **Goat anti-Rabbit Alexa Fluor 488** | A-11008 | Thermo Fisher Scientific | **IMF**: 1:200 |
| **Goat anti-Mouse IgG Alexa Fluor 488** | A-11001 | Thermo Fisher Scientific | **IMF**: 1:200 |

#### **Supplementary Table S3-** List of primers used in this study

Underlined text indicate restriction site sequences and bold text mutated bases. SDM: site directed mutagenesis

| Name | Sequence 5’-3’ | Purpose |
|---|---|---|
| **hnRNP U_NTD_Fwd** | GAAGATGAATTCATGAGTTCCTCGCCTGT | Cloning |
| **hnRNP U_NTD_Rev** | ATGCGAGTCGACTCAATACTTGTTCTCTTC | Cloning |
| **hnRNP U_SPRY_Fwd** | ATTCCGGAATTCAGCAGAGCCAAATCTCC | Cloning |
| **hnRNP U_SPRY_Rev** | TGCCAAGTCGACTCACTTTTCCTTCTGACC | Cloning |
| **hnRNP U_CTD_Fwd** | TGCCAAGAATTCCCATATTTTCCAATACC | Cloning |
| **hnRNP U_CTD_Rev** | TTCCGGTCGACTCAATAATATCCTTGG | Cloning |
| **GRP1-PH K273A fwd** | 5’ GAA GGC TGG CTG CTG **GC**G CTG GGG GGT CG | SDM |
| **GRP1-PH K273A rev** | 5’ CG ACC CCC CAG C**GC** CAG CAG CCA GCC TTC | SDM |

#### Supplementary Figure S1

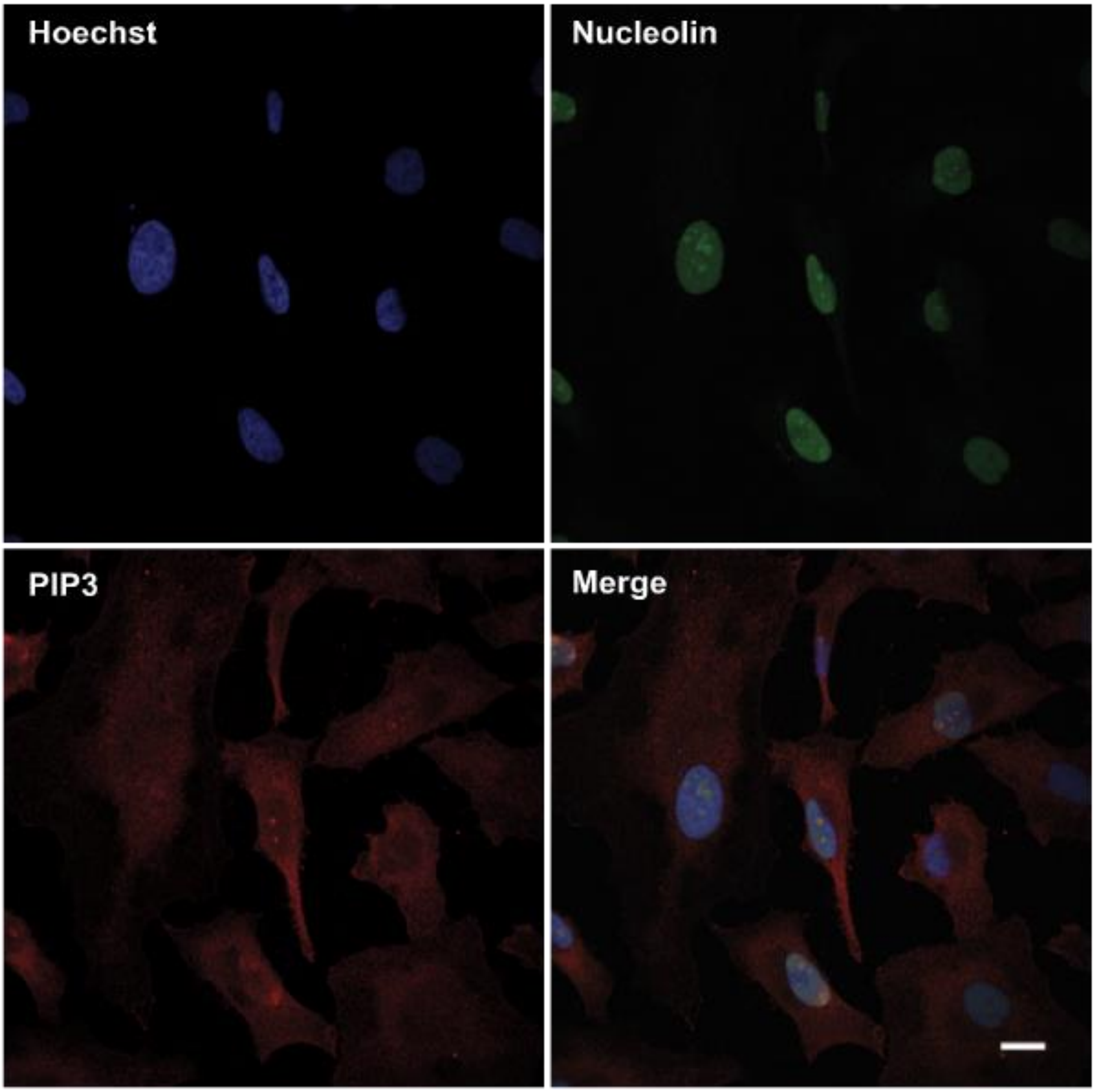


**Supplementary Figure S1. Overview of PtdIns(3,4,5)*P*3 nucleolar localisation in HeLa cells**

Co-immunostaining of HeLa cells was performed with the indicated antibodies and imaged by confocal microscopy. PIP3: PtdIns(3,4,5)$P_3$; NPM1: nucleophosmin; Scale bar represents 10 µm.

#### Supplementary Figure S2

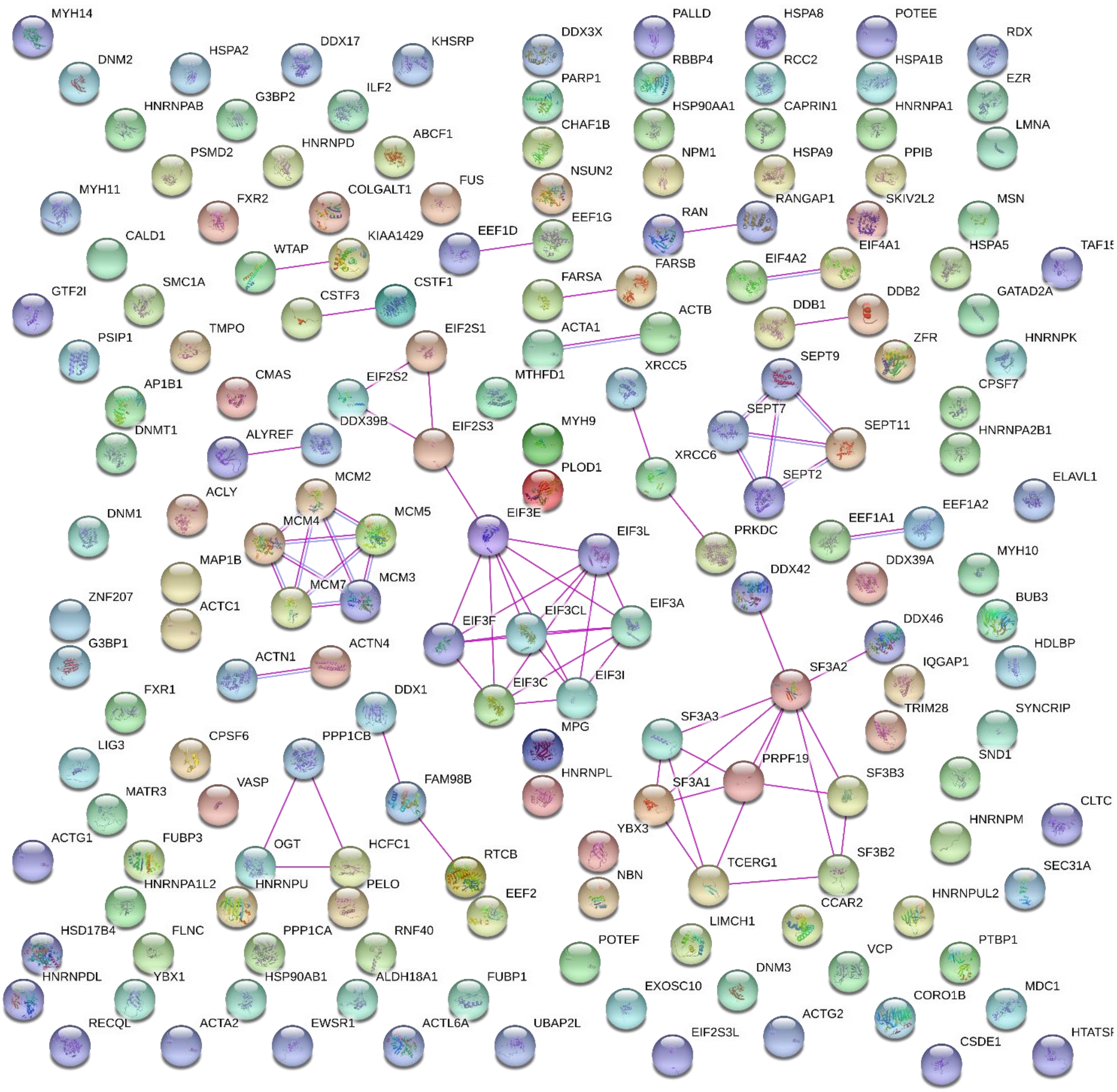


**Supplementary Figure S2. Overview of potential protein-protein interaction of the PtdIns(3,4,5)*P*$_3$ binding proteins identified in this study.**

STRING version 10.5 ([https://st ring-db.org/](https://string-db.org/) (1)) analysis of the PtdIns(3,4,5)*P*$_3$ binding proteins using data settings enabling only experimental evidence to be considered and a confidence score > 0.9. Protein-protein complexes are shown with connecting pink and/or purple lines. Amongst these interaction complexes, a few proteins harbour a K/R motif, *i.e.* DDX39B, MCM3, SEPT7, ZFR, SF3A1, DDX42 and RAN.

#### Supplementary Figure S3

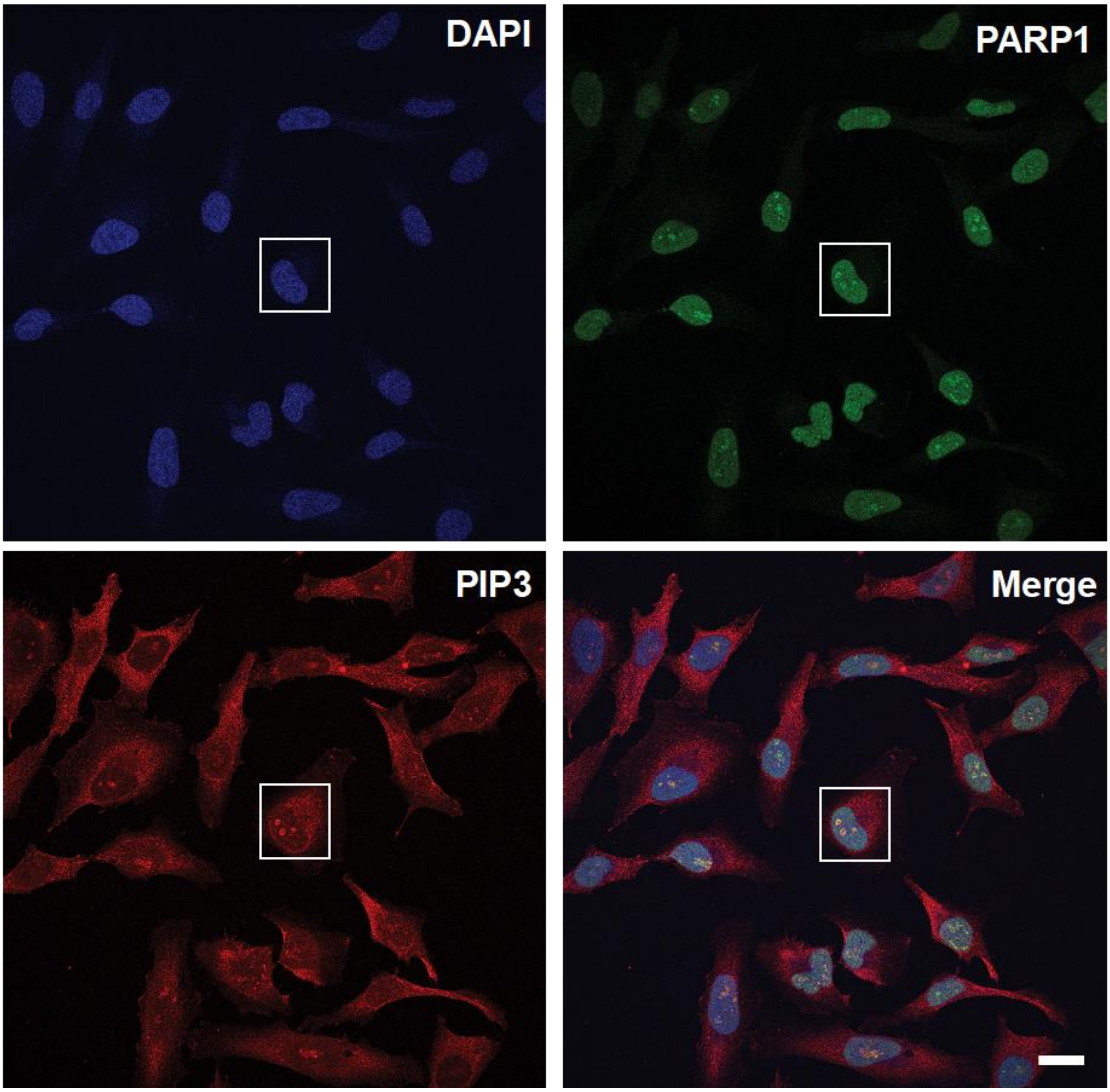


**Supplementary Figure S3. Overview of the PARP1-PtdIns(3,4,5)$P_3$ nucleolar colocalisation in actively growing HeLa cells**

Asynchronous HeLa cells were co-stained with anti-PARP1 and PtdIns(3,4,5)$P_3$ antibodies and imaged by confocal microscopy. Scale bar represents 10 µm.
